# Self-assembling short immunostimulatory duplex RNAs with broad spectrum antiviral activity

**DOI:** 10.1101/2021.11.19.469183

**Authors:** Longlong Si, Haiqing Bai, Crystal Yuri Oh, Tian Zhang, Fan Hong, Amanda Jiang, Yongxin Ye, Tristan X. Jordan, James Logue, Marisa McGrath, Chaitra Belgur, Atiq Nurani, Wuji Cao, Rachelle Prantil-Baun, Steven P Gygi, Rani K. Powers, Matthew Frieman, Benjamin R. tenOever, Donald E. Ingber

## Abstract

The current COVID-19 pandemic highlights the need for broad-spectrum antiviral therapeutics. Here we describe a new class of self-assembling immunostimulatory short duplex RNAs that potently induce production of type I and type III interferon (IFN-I and IFN-III), in a wide range of human cell types. These RNAs require a minimum of 20 base pairs, lack any sequence or structural characteristics of known immunostimulatory RNAs, and instead require a unique conserved sequence motif (sense strand: 5’-C, antisense strand: 3’-GGG) that mediates end-to-end dimer self-assembly of these RNAs by Hoogsteen G-G base-pairing. The presence of terminal hydroxyl or monophosphate groups, blunt or overhanging ends, or terminal RNA or DNA bases did not affect their ability to induce IFN. Unlike previously described immunostimulatory siRNAs, their activity is independent of TLR7/8, but requires the RIG-I/IRF3 pathway that induces a more restricted antiviral response with a lower proinflammatory signature compared with poly(I:C). Immune stimulation mediated by these duplex RNAs results in broad spectrum inhibition of infections by many respiratory viruses with pandemic potential, including SARS-CoV-2, SARS-CoV, MERS-CoV, and influenza A, as well as the common cold virus HCoV-NL63 in both cell lines and human Lung Chips that mimic organ-level lung pathophysiology. These short dsRNAs can be manufactured easily, and thus potentially could be harnessed to produce broad-spectrum antiviral therapeutics at low cost.

## INTRODUCTION

Recognition of duplex RNAs by cellular RNA sensors plays a central role in host response to infections by initiating signaling cascades that induce secretion of interferon (IFN) and subsequent upregulation of hundreds of interferon-stimulated genes (ISGs). This pathway therefore also serves as a potent point of therapeutic intervention in a broad range of viral diseases. Duplex RNAs with various structural features have been identified that are recognized by the three cellular RNA sensors that are responsible for this innate immune response (1). One of these, toll-like receptor 3 (TLR3), is located on the cell membrane and the endosomal membrane, while the other two-retinoic acid inducible gene I (RIG-I) and melanoma differentiation associated gene 5 (MDA5)-are located in the cytosol. Long forms of duplex RNA are recognized by these sensors based on their length (i.e., independently of the structure of their 5’ ends) with TLR3 recognizing duplex RNAs >35 bp and MDA5 sensing duplex RNAs >300 bp (2). Past reports have revealed that a short stretch of duplex RNA (>19 bp) can be recognized by RIG-I, but only if a triphosphate or a diphosphate is present at its 5′ end and if the end is blunt with no overhangs (1,3–5).

Duplex RNA-mediated innate immune stimulation is a two-edged sword. For example, in the case of respiratory infections, such as those caused by pandemic viruses (e.g., SARS-CoV-2, SARS-CoV, MERS-CoV, and influenza virus), RNA-mediated activation of this innate immune response provides the first line of host defense against the invading pathogen. However, on the other hand, the use of duplex RNAs for RNA interference (RNAi) approaches can result in undesired immunological off-target effects and misinterpretation of experimental results (6–12). Thus, gaining greater insight into the mechanism by which cells sense and respond to duplex RNAs could have broad impact in biology and medicine.

In this study, we serendipitously discovered a class of new immunostimulatory RNAs while using >200 small interfering RNAs (siRNAs) to identify influenza infection-associated host genes in human lung epithelial cells. These short duplex RNAs potently induce type I and type III interferons (IFN-I/III) in a wide type of cells, but lack any sequence or structure characteristics of known immunostimulatory RNAs. Systematic mechanistic analysis revealed that these immunostimulatory RNAs specifically activate the RIG-I/IRF3 pathway by binding directly to RIG-I, and that this only occurs when these short RNAs have a conserved overhanging sequence motif (sense strand: 5’-C, antisense strand: 3’-GGG) and a minimum length of 20 bases. Interestingly, the conserved overhanging motif is responsible for the self-assembly of end-to-end RNA dimers through Hoogsteen G-G base pairing. In addition, these immunostimulatory RNAs appear to be novel in that they are capable of inducing IFN production regardless of whether they have blunt or overhanging ends, terminal hydroxyl or monophosphate groups, RNA base-or DNA base-ends, in contrast to previously described immunostimulatory RNAs that require 5’-di or triphosphates to activate cellular RNA sensors (3, 4). The RNA-mediated IFN-I/III production resulted in significant inhibition of infections by multiple human respiratory viruses, including influenza viruses and SARS-CoV-2 in established cell lines and in human Lung Airway and Alveolus Chips that have been previously shown to recapitulate human lung pathophysiology (13–15). These findings also should facilitate the development of siRNAs that avoid undesired immune activation and may pave the way for the development of a new class of RNA therapeutics for the prevention and treatment of respiratory virus infections.

## RESULTS

### Discovery of IFN-I pathway-activating immunostimulatory RNAs

While using >200 siRNAs to identify host genes that mediate human A549 lung epithelial cell responses to influenza A/WSN/33 (H1N1) infection, we found that transfection of two siRNAs (RNA-1 and RNA-2) inhibited H1N1 replication by more than 90% (Fig. 1A). To explore the mechanism of action of these siRNAs, we profiled the transcriptome and proteome of A549 cells transfected with RNA-1 (Fig. 1B) and RNA-2 (Fig. S1), which respectively target the long non-coding RNAs (lncRNAs) DGCR5 and LINC00261, and a scrambled siRNA was used as a control. RNA-seq analysis showed that RNA-1 upregulates the expression of 21 genes by more than 2-fold (*p* value threshold of 0.01) (Fig. 1B **left** and Fig. S2A **left**). Gene Oncology (GO) enrichment analysis revealed that these genes are involved in IFN-I signaling pathway and host defense response to viral infections (Fig. 1B **left**), including MX1, OASL, IFIT1, and ISG15 (Fig. S2A **left**). In parallel, Tandem Mass Tag Mass Spectrometry (TMT Mass Spec) quantification demonstrated upregulation of 73 proteins by more than 4-fold (*p* value threshold of 0.01), including IL4I1, TNFSF10, XAF1, IFI6, and IFIT3 (Fig. 1B **right** and Fig. S2B). GO enrichment analysis of these upregulated proteins also confirmed an association between treatment of RNA-1 and induction of the IFN-I pathway (Fig. 1B **right** and Fig. S3A). Quantitative reverse transcription polymerase chain reaction (qRT-PCR) assay independently validated that RNA-1 preferentially activates the IFN-I pathway relative to the Type II IFN pathways (Fig. S3B), with IFN-β being induced to much higher levels (>1,000-fold) compared to IFN-α (Fig. 1C). This potent induction of IFN-β by RNA-1 was verified at the protein level using enzyme-linked immunosorbent assay (ELISA) (Fig. S4), and similar patterns of gene and protein expression were also observed for RNA-2 (Fig. 1C and Figs. S1 **to** S2).

**Figure 1.**
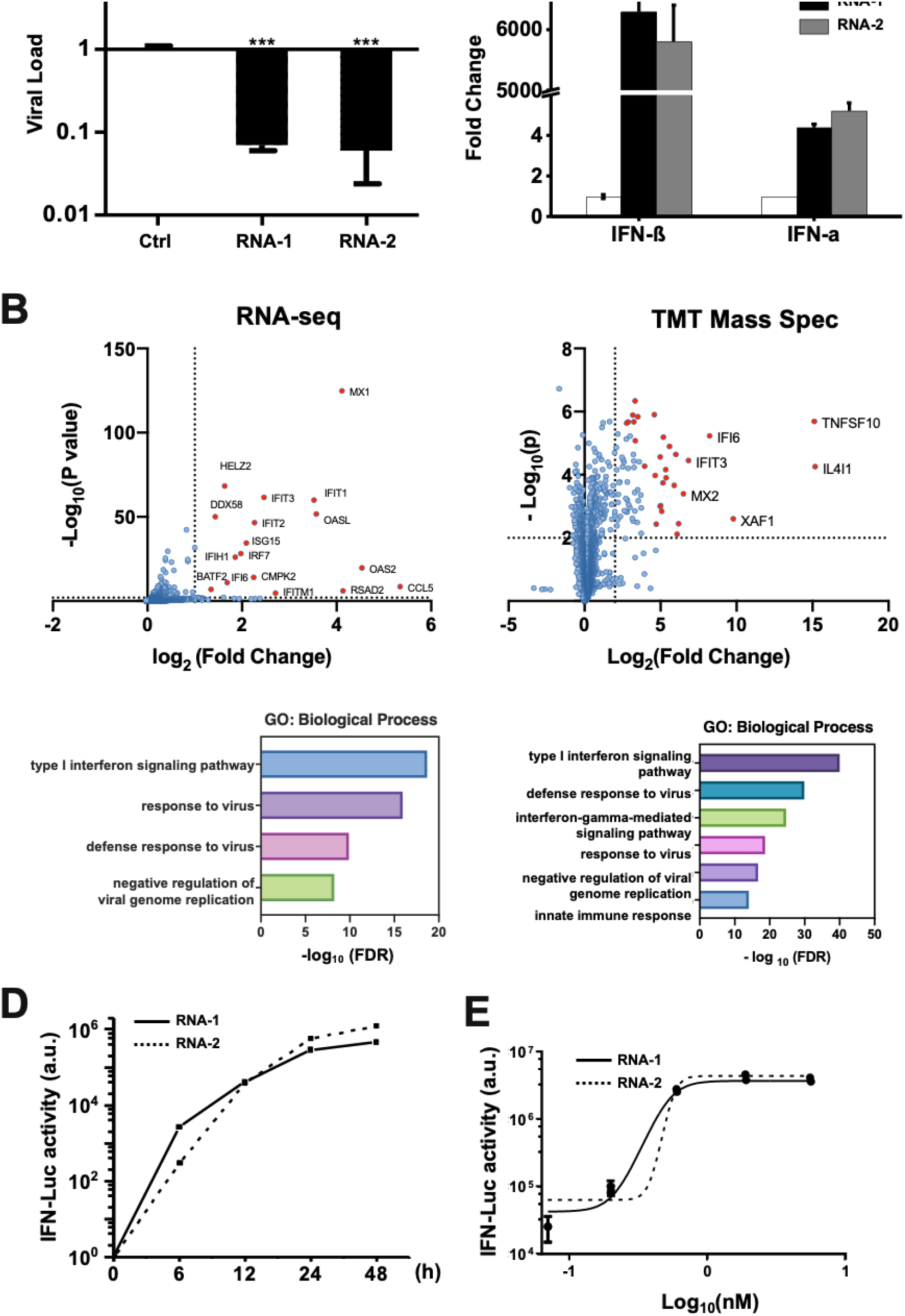
Discovery of new immunostimulatory RNAs. (**A**) A549 cells were transfected with RNA-1, RNA-2, or a scrambled duplex RNA control, and infected with influenza A/WSN/33 (H1N1) virus (MOI=0.01) 24 hours later. Titers of progeny viruses in medium supernatants collected at 48 h post-infection were determined by quantifying plaque forming units (PFUs); data are shown as % viral infection measured in the cells treated with the control RNA (Data shown are mean ± standard deviation; N =3; ***, *p* < 0.001). (**B**) A549 cells were transfected with RNA-1 or a scrambled dsRNA control, collected at 48 h, and analyzed by RNA-seq (left) or TMT Mass Spec (right). Differentially expressed genes (DEGs) from RNA-seq or proteins from TMT Mass Spec are shown in volcano plots (top) and results of GO Enrichment analysis performed for the DEGs are shown at the bottom (N = 3). (**C**) qPCR analysis of cellular IFN-β and IFN-α RNA levels at 48 h after A549 cells were transfected with RNA-1, RNA-2, or scrambled dsRNA control (N = 3). (**D**) RNA-mediated production kinetics of IFN production in wild-type A549-Dual cells that were transfected with RNA-1, RNA-2, or scramble RNA control measured using a Quanti-Luc assay. OD values from cells transfected with the scrambled RNA control were subtracted as background (N = 6). (**E**) Dose-dependent induction of IFN by RNA-1 and -2 in A549-Dual cells compared to scrambled RNA control measured at 48 h post-transfection (control OD values were subtracted as background; N = 6).

Interestingly, when we carried out studies with additional siRNAs to further validate the function of the lncRNAs they target, we found that knockdown of DGCR5 or LINC00261 by these other siRNAs did not induce IFN production. This was surprising because since the inception of RNA interference technology, short duplex (double stranded) siRNAs have been known to induce IFN-I (7, 9) and thus subsequent design of these molecules, including the ones used in our study, were optimized to avoid this action and potential immunomodulatory side effects (16). siRNAs synthesized by phage polymerase that have a 5’-triphosphate end can trigger potent induction of IFN-α and -β (7), and siRNAs containing 9 nucleotides (5’-GUCCUUCAA-3’) at the 3’ end can induce IFN-α through TLR-7 (8). Notably, RNAs with a 5’-diphosphate end can induce IFN-I as well (17), but our synthetic duplex RNAs do not have any of these sequence or structural properties. Thus, our data suggested that the two specific RNAs we found to be potent IFN-I/III inducers (RNA-1 and RNA-2) may represent new immunostimulatory RNAs.

To explore this further, we assessed IFN production induced by the two putative immunostimulatory RNAs using an A549-Dual^TM^ IFN reporter cell line, which stably expresses luciferase genes driven by promoters containing IFN-stimulated response elements (18). These studies revealed that both RNA-1 and -2 induce IFN production beginning as early as 6 hours post transfection, consistent with IFN-I/III being an early-response gene in innate immunity, and high levels of IFN expression were sustained for at least 24 to 48 hours (Fig. 1D). We also observed dose-dependent induction of IFN production by these duplex RNAs over the nM range (Fig. 1E). In addition, we observed similar effects when we tested RNA-3, which was originally designed as a siRNA to knockdown another lncRNA, LINC00885 (Fig. 2, Table 1). Notably, all three immunostimulatory dsRNAs that specifically upregulate strong IFN-I/III responses with high efficiency share a common motif (sense strand: 5’-C, antisense strand: 3’-GGG).

**Figure 2.**
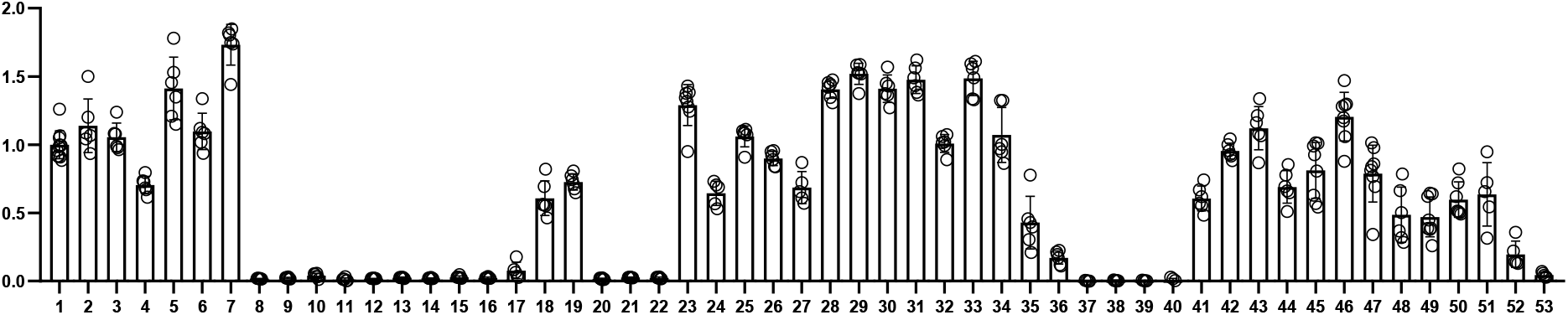
Comparison of the immunostimulatory activities of different RNAs. A549-Dual cells were transfected with indicated duplex RNAs for 48 h, and then activation of the IFN pathway was measured by quantifying luciferase reporter activity. The immunostimulatory activity of RNA-1 was set as 1 (N = 6).

**Table 1.**
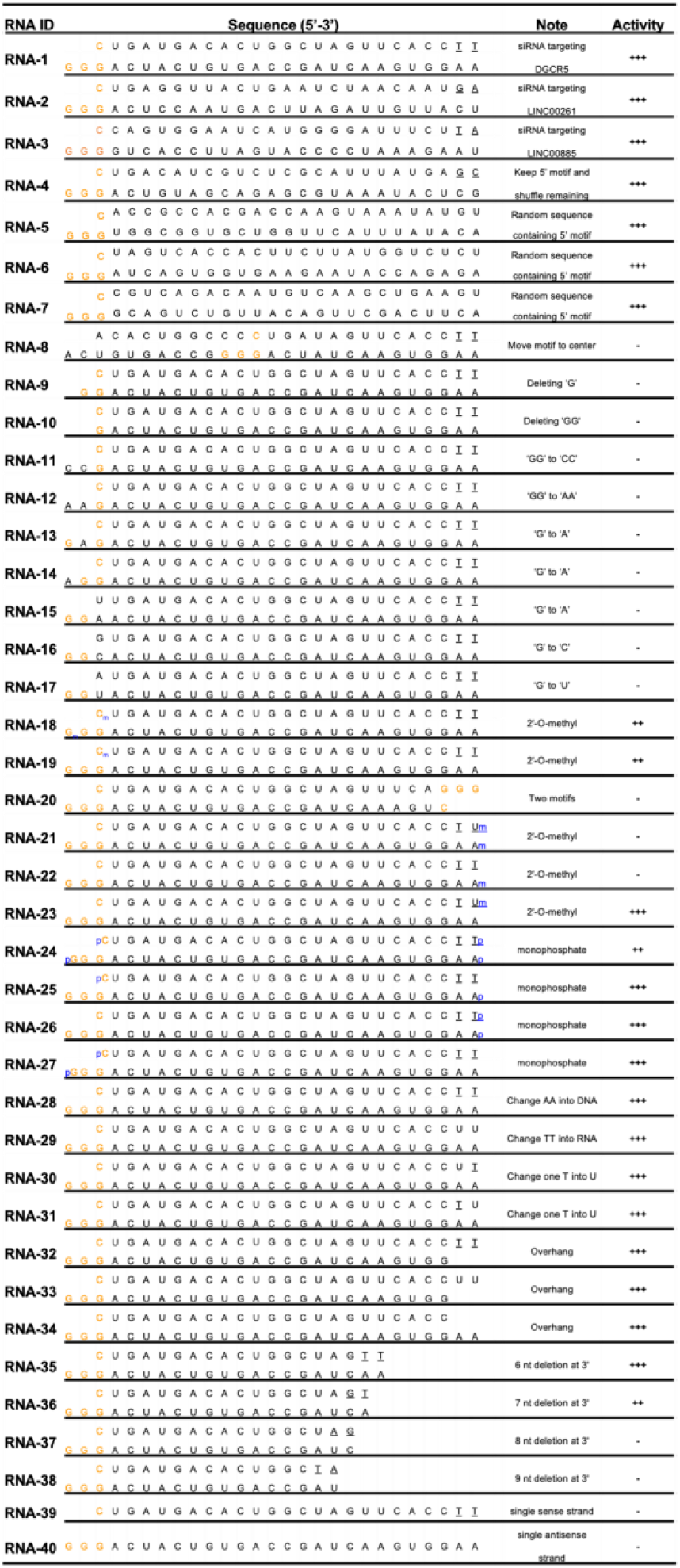
Oligonucleotides of RNA monomers. The sense strand (left 5’ end-right 3’ end) is positioned on top, while the antisense strand (left 3’ end-right 5’ end) is below. If not indicated otherwise, both 5’ and 3’ ends of sense and antisense contain terminal hydroxyl groups. The underlined bases indicate DNA bases; p, monophosphate group; m, N_1_-2-O-methyl group. +++, high activity; ++. Middle activity; +, low activity; -, no activity.

### These short duplex RNAs bind directly to RIG-I

Transcription factor interferon regulatory factor 3 (IRF3) and 7 (IRF7) play vital roles in IFN-I production (19, 20). Using IRF3 knockout (KO) and IRF7 KO cells, we found that loss of IRF3, but not IRF7, completely abolished the ability of RNA-1 to induce IFN-β (Fig. 3A) and downstream ISGs, including STAT1, IL4L1, TRAIL, and IFI6 (Fig. S5). IRF3 is the master and primary transcriptional activator of IFN-I and its induction of IFN-I involves a cascade of events, including IRF3 phosphorylation, dimerization, and nuclear translocation (21, 22). To alleviate potential interference from host gene knockdown by RNA-1 that was developed as an siRNA, we performed further mechanistic studies using RNA-4, which contains the common motif of RNA-1, -2, and -3 that we hypothesized and proved to be involved in the immunostimulatory activity, but does not target (silence) any host genes because its other nucleotides were randomized (Fig. 2, Table 1). Although RNA-4 had no effect on IRF3 mRNA or total protein levels (Fig. 3B,C), it increased IRF3 phosphorylation (Fig. 3C), which is essential for its transcriptional activity (19) and subsequent translocation to the nucleus (Fig. 3D), where IRF3 acts as transcription factor that induces IFN-I expression (21, 22).

**Figure 3.**
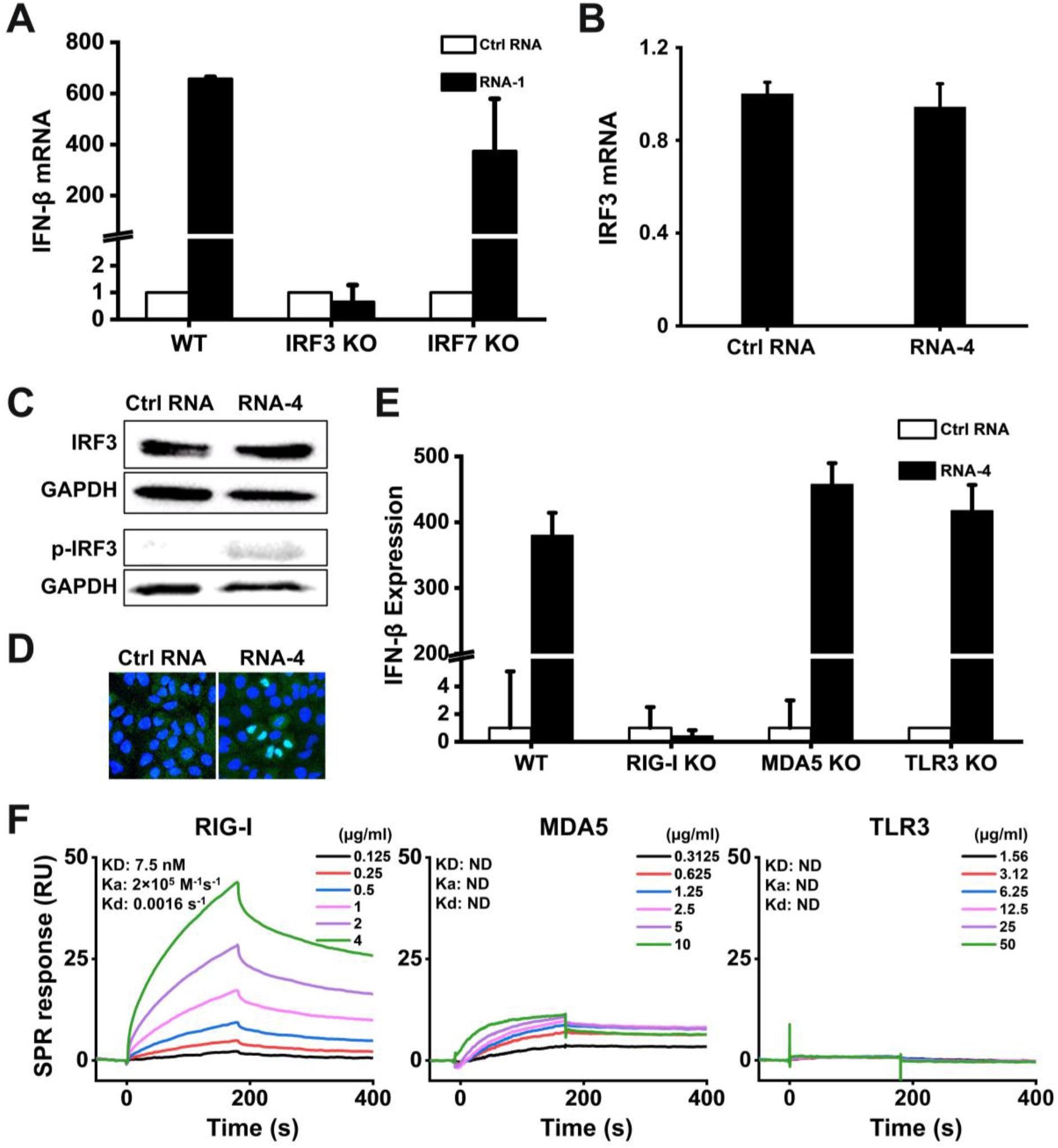
Immunostimulatory RNAs induce IFN-I production through RIG-I-IRF3 pathway. (**A**) Wild-type (WT) HAP1 cells, IRF3 knockout HAP1 cells, or IRF7 knockout HAP1 cells were transfected with RNA-1 or scrambled RNA control for 48 h, and IFN-β mRNA levels were quantified by qPCR. Data are shown as fold change relative to the scrambled RNA control (N = 3). Note that IRF3 knockdown completely abolished the IFN-β response. (**B**) IRF3 mRNA levels measured in A549 cells transfected with immunostimulatory RNA-4 or a scrambled RNA control, as determined by qPCR and 48 h post-transfection (data are shown as fold change relative to the control RNA; N = 3). (**C**) Total IRF3 protein and phosphorylated IRF3 detected in A549 cells transfected with RNA-4 or scrambled RNA control at 48 h post transfection as detected by Western blot analysis (GAPDH was used as a loading control). (**D**) Immunofluorescence micrographs showing the distribution of phosphorylated IRF3 in A549 cells transfected with RNA-4 or scrambled RNA control at 48 h post transfection (Green, phosphorylated IRF3; blue, DAPI-stained nuclei; arrowheads, nuclei expressing phosphorylated IRF3). (**E**) Wild-type (WT) A549-Dual cells, RIG-I knockout A549-Dual cells, MDA5 knockout A549-Dual cells, or TLR3 knockout A549 cells were transfected with immunostimulatory RNA-4 or a scrambles RNA control and 48 h later, IFN-β expression levels were quantified using the Quanti-Luc assay or qPCR (data are shown as fold change relative to the scrambled RNA control; N = 6). Note that RIG-I knockout abolished the ability of the immunostimulatory RNAs to induce IFN-β. (**F**) SPR characterization of the binding affinity between cellular RNA sensors (RIG-I, MDA5, and TLR3) and RNA-1, which were immobilized on a streptavidin (SA) sensor chip. Equilibrium dissociation constant (KD), association rate constant (Ka), and dissociation rate constant (Kd) are labeled on the graphs.

RIG-I, MDA5, and TLR3 are the main sensors upstream of IRF3 that recognize RNA (23). To investigate which of them detect the immunostimulatory short duplex RNAs, we quantified RNA-mediated production of IFN-I in RIG-I, MDA5, or TLR3 KO cells. Knockout of RIG-I completely suppressed the ability of RNA-4 (Fig. 3E) as well as RNA-1 and -2 (Fig. S6) to induce IFN-I, whereas loss of MDA5 or TLR3 had no effect on RNA-mediated IFN-I production (Fig. 3E and Fig. S6). Importantly, surface plasmon resonance (SPR) analysis revealed that RNA-1 interacts directly with the RIG-I cellular RNA sensor, rather than MDA5 or TLR3 (Fig. 3F). In addition, knockout or overexpression of other RNA sensors, such as TLR7 or TLR8, did not affect the ability of these duplex RNAs to induce IFN production (Fig. S7). Thus, these short duplex RNAs stimulate IFN-I production specifically via the RIG-I/IRF3 pathway.

### Overhanging GGG motif mediates IFN activation via duplex RNA dimerization

The active RNAs-1, -2, and -3 are chemically synthesized 27-mer RNA duplexes that include terminal hydroxyl groups, 2 DNA bases at the 3’ end of sense strands, and 2-base overhangs at the 3’ end of antisense strands (Table 1). Importantly, their sequence and structure features do not conform to any characteristics of existing immunostimulatory RNA molecules (Table S1), suggesting that previously unknown elements must be responsible for this immunostimulatory activity. Remarkably, even though they were designed to target different host genes, sequence alignment revealed that RNA-1, -2, and -3 contained one identical motif at their 5’ ends (sense strand: C, antisense strand: 3’-GGG-5’) (Table 1). Because all the three RNAs were potent inducers of IFN, we hypothesized that this common motif may mediate their immunostimulatory activities.

To test this hypothesis, we systematically investigated IFN production induced by different sequence variants of RNA-1 (Table 1) using the IFN reporter-expressing cell line. Maintaining the common motif while shuffling remaining nucleotides or replacing them with a random sequence (RNA-4 or RNA-5, -6, and -7, respectively, *vs.* RNA-1, -2, and -3) did not affect the immunostimulatory activity of the duplex RNA (Fig. 2 and Table 1). However, moving the motif from 5’ GGG end to the middle region completely abolished the RNA’s immunostimulatory activity (RNA-8 *vs.* RNA-1) (Fig. 2 and Table 1). Furthermore, the immunostimulatory activity was completely eliminated by any changes, including deletion or substitution, at the common motif (RNA-9, -10, -11, -12, -13, -14, -15, -16, -17 *vs.* RNA-1) (Fig. 2 and Table 1). These data indicate that the common terminal 5’ GGG motif is necessary for IFNI/III induction, and that this effect is sensitive to alterations in its position and sequence.

To determine whether this shared motif mediates binding to RIG-I, we evaluated the immunostimulatory activity of duplex RNAs bearing an N_1_-2’O-methyl group, which has been shown to block RIG-I activation by RNA when the modification occurs at the 5’-terminus (24). Surprisingly, the N_1_-2’O-methylation of the 5’-end of sense strand or 3’-end of antisense strand (RNA-18 and -19) or both simultaneously in the same duplex RNA (RNA-20) did not block RIG-I activation by RNA-1 (Fig. 2 and Table 1). In contrast, N_1_-2’O-methylation of the 5’-end of the antisense strand, but not the 3’-end of the sense strand, completely blocked RIG-I activation by RNA (RNA-21, -22, and -23 *vs.* RNA-1) (Fig. 2 and Table 1), indicating that RNA-1 binds to RIG-I via the 5’GGG-end of its antisense strand.

Given the critical role and high conservation of the common motif in this form of duplex RNA-mediated immunostimulation, we also explored whether this common motif could mediate the formation of higher order structure of duplex RNA via an intramolecular G-quadruple, a secondary structure that is held together by non-canonical G-G Hoogsteen base pairing (25). Interestingly, native gel electrophoresis revealed the formation of an RNA-1 dimer, while no dimer was detected when the GG overhang was replaced with AA bases (RNA-12 *vs.* RNA-1) (Fig. 4A). These data suggest that the common motif (sense strand: 5’-C, antisense strand: 3’-GGG-5’) mediates formation of an end-to-end RNA-1 dimer via Hoogsteen G-G base pairing (25), which doubles the length of the dsRNA, thereby promoting efficient binding to RIG-I via the exposed 5’ antisense strand ends of each RNA and subsequently inducing IFN production (Fig. 4B). This possibility was verified by synthesizing RNA-1 tail-to-tail dimer mimics (RNA-41 and -42) that have similar lengths and sequences and also exhibited potent immunostimulatory activity (Fig. 2, Table 2).

**Figure 4.**
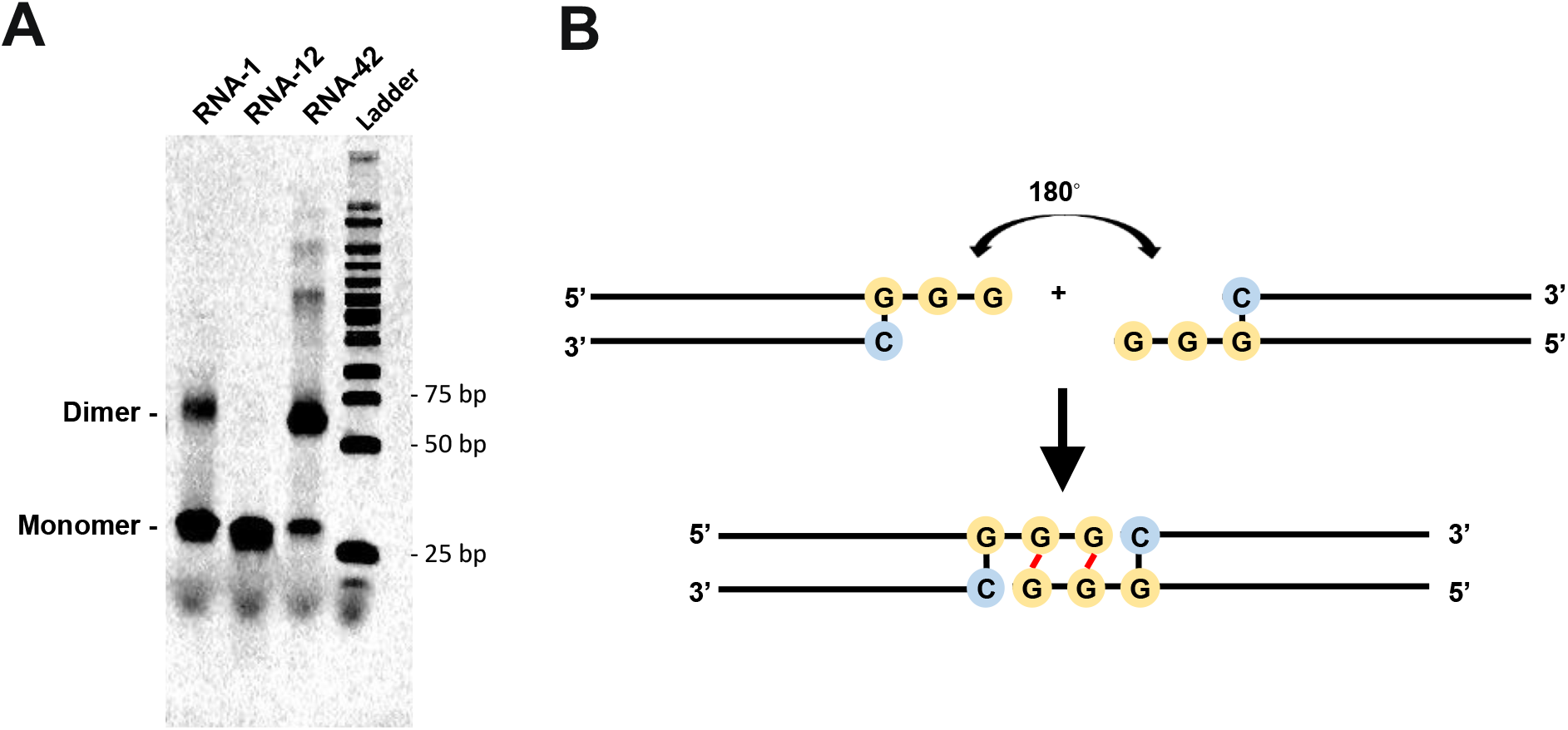
The common motif mediates the formation of duplex RNA dimers via intramolecular G-quadruplex formed by GG overhang. (**A**) The image of native gel electrophoresis showing the formation of RNA-1 dimer. 1 uL of 10 uM RNA samples were loaded. RNA-12 and RNA-42 were used as negative and positive control, respectively. (**B**) The diagram showing the structure of ‘end-to-end’ RNA-1 dimer due to terminal G-G Hoogsteen paring.

**Table 2.**
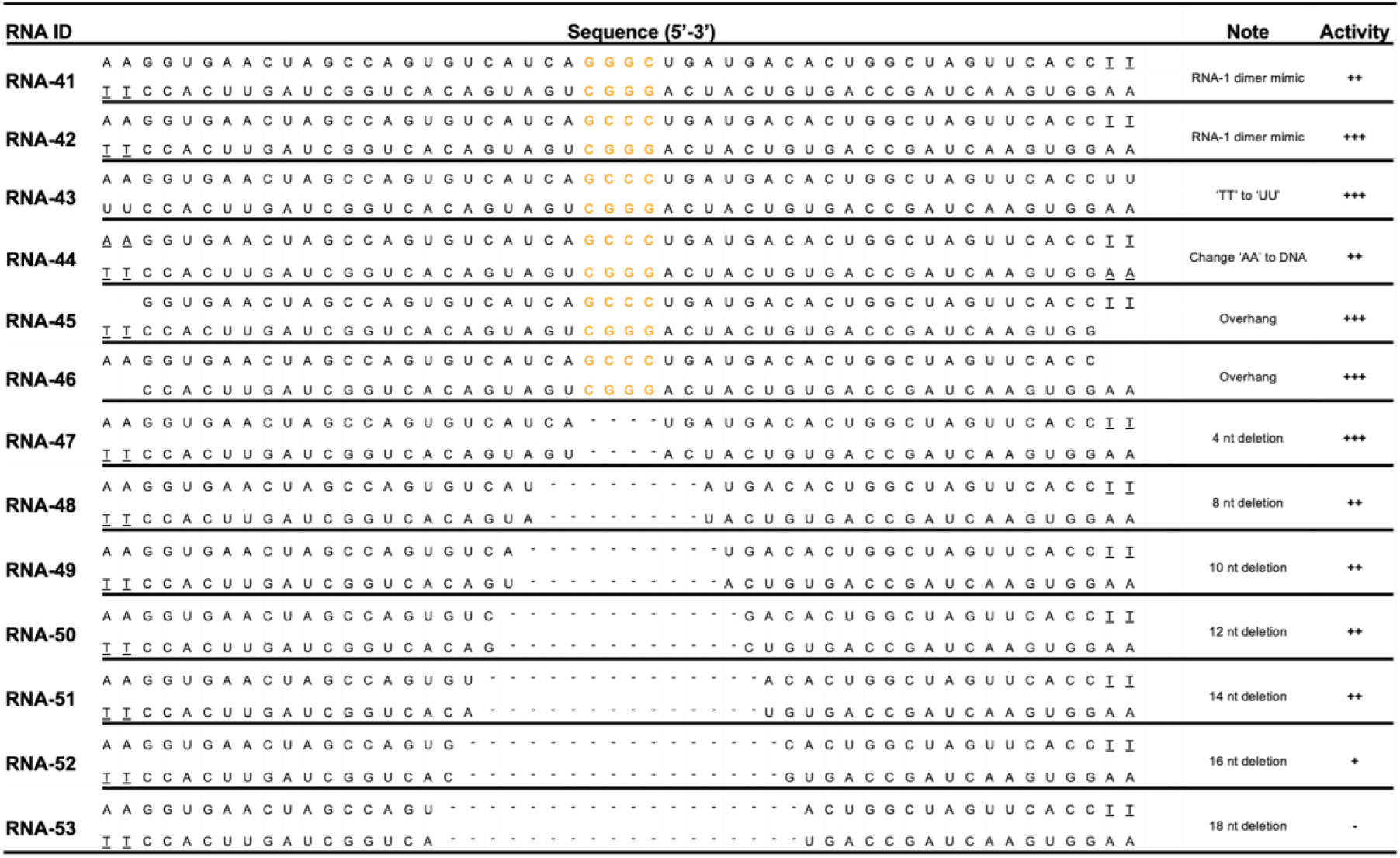
Oligonucleotides of RNA-1 dimer mimics. The sense strand (left 5’ end-right 3’ end) is positioned on top, while the antisense strand (left 3’ end-right 5’ end) is below. If not indicated otherwise, both 5’ and 3’ ends of sense and antisense contain terminal hydroxyl groups. The underlined bases indicate DNA bases; p, monophosphate group; m, N_1_-2-O-methyl group. +++, high activity; ++. Middle activity; +, low activity; -, no activity.

As chemically synthesized RNAs contain terminal hydroxyl groups, we tested if adding a monophosphate at these sites affects the IFN-inducing activity. This is important to investigate because host RNAs contain a 5’-monophosphate, which has been reported to suppress RIG-I recognition (3). However, we found that RNA-1 containing terminal monophosphates exhibited immunostimulatory activity to a similar level as RNA-1 containing a hydroxyl groups (RNA-24, -25, -26, -27 *vs.* RNA-1) (Fig. 2, Table 1), suggesting that a terminal monophosphate in these short duplex RNAs is neither required, nor does it interfere with, their immunostimulatory activity.

As our dsRNAs contain 2 DNA bases at the 3’ end of their sense strand, we also tested if the types of nucleosides affect the IFN-inducing activity. Interestingly, the duplex RNAs exhibited comparable immunostimulatory activity to RNA-1 regardless of whether DNA bases or RNA bases are inserted at the 3’ end of the sense strand and/or 5’ end of the antisense strand (RNA-28, -29, -30, -31 *vs.* RNA-1) (Fig. 2, Table 1). This was further verified by synthesizing duplex RNA dimer mimics (RNA-43 and -44 *vs.* RNA-42) that contain terminal DNA or RNA bases, which exhibited similar immunostimulatory activity to RNA-42 that contains 2 DNA bases at the 3’ ends of sense and antisense strands (Fig. 2, Table 2).

We then tested if introduction of an overhang affects the IFN-inducing activity because previous reports revealed that RIG-I can be activated by blunt duplex RNAs, and that almost any type of 5’ or 3’ overhang can prevent RIG-I binding and eliminate signaling (4). However, we found that the overhang did not affect the IFN-inducing activity of our duplex RNAs (RNA-32, -33, and -34 *vs.* RNA-1) (Fig. 2, Table 1). This was also verified in studies with duplex RNA mimics (RNA-45 and -46 *vs.* RNA-42) that contain terminal overhangs, which induced IFN production to a similar level as RNA-42 containing blunt ends (Fig. 2**, Table2**).

Finally, we analyzed the effects of RNA length on IFN production by gradually trimming bases from the 3’ end of RNA-1. Removal of increasing numbers of bases resulted in a gradual decrease in immunostimulatory activity (RNA-35 and -36 vs. RNA-1) with complete loss of activity when 8 bases or more were removed from the 3’ end of RNA-1 (RNA-37 and -38) (Fig. 2, Table 1). Therefore, the minimal length of this new form of immunostimulatory RNA required for IFN induction is 20 bases on the antisense strand that can result in the formation of a RNA dimer containing ∼38 bases via Hoogstein base pairing of their 5’GG ends. This is consistent with data obtained with duplex RNA tail-to-tail dimer mimics (RNA-47, -48, -49, -50, -51, -52, and -53 *vs.* RNA-41 and -42) where the minimal length of the duplex RNA dimer required for IFN induction was found to be 36 bases (Fig. 2, Table 2). And in a final control experiment we found that neither the single sense strand nor the single antisense strand of RNA-1 alone is sufficient to induce IFN production (RNA-39 and -40) (Fig. 2, Table 1), indicating that the double stranded RNA structure is absolutely required for its immunostimulatory activity.

Finally, given that the overhanging motif (sense strand: C; antisense strand: 3’-GGG-5’) is also found in the termini of many siRNAs that can be immunostimulatory, we evaluated its frequency in both human mRNAs and lncRNAs. Genome-wide sequence analysis revealed that the ‘CCC’ motif is abundant in both mRNAs and lncRNAs sequences: 99.96 % of human mRNAs contain ‘CCC’ with an average distance of 75.45 bp between adjacent motifs and 98.08 % of human lncRNAs contain ‘CCC’ with an average distance of 75.93 bp between adjacent motifs (Fig. S8). Thus, this indicates that the ‘GGG’ motif that mediates short duplex RNA dimerization should be avoided when an siRNA’s immunostimulatory effect is undesired. ***Self-assembling dsRNAs induce less proinflammatory genes than poly(I:C)***

Polyinosinic:polycytidylic acid [poly(I:C)] is an immunostimulant used to simulate viral infections, which interacts with multiple pattern recognition receptors, including toll-like receptor 3 (TLR3), RIG-I, and MDA5. To compare the immunostimulatory landscape induced by RNA-1 with poly(I:C), we performed bulk RNA-seq analysis of A549 cells transfected with the sample amounts of scrambled dsRNA as control, RNA-1, or poly(I:C) for 48 hours. Principle-component analysis shows that RNA-1 and poly(I:C) induce distinct transcriptomic changes (Fig. 5A). Similar to earlier results (Fig. 1B), RNA-1 upregulated many genes that are involved in antiviral IFN response genes, such as *MX1, OASL, IRF7, IFIT1* (Fig. 5B). In contrast, poly(I:C) induces much broader changes in gene expression: 302 genes have decreased expression while only 2 decrease when treated with RNA-1 (Fig. 5C). A heat map also shows that many proinflammatory cytokines and chemokines, such as *CXCL11*, *TNF*, *CCL2, IL1A*, have much higher expression in cells transfected with poly(I:C) (Fig. 5D). In addition, a number of genes involved in ion transport and cell adhesion are decreased by poly(I:C) but not by RNA-1. Notably, many of these genes (*MYO1A, NEB, ADH6, H19, ELN*, etc.) were also down-regulated in SARS-CoV-2 infection (26). These results indicate that, when compared to poly(I:C), our dsRNAs induce a more targeted antiviral response and a lower level of tissue-damaging proinflammatory responses, while having no effect on critical biological processes, such as ion transport and cell adhesion, which should make them more suitable for antiviral therapeutic applications.

**Figure 5.**
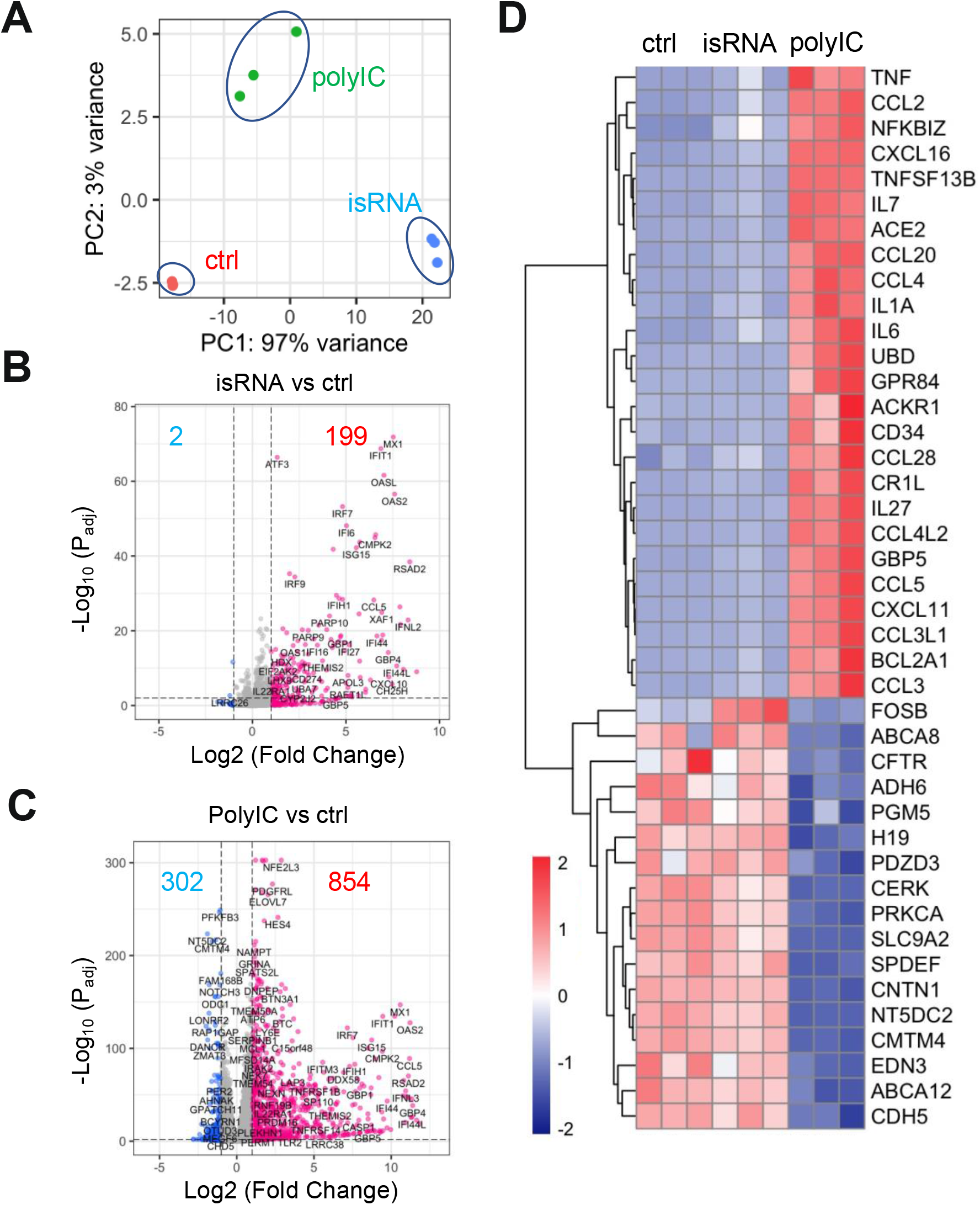
Immunostimulatory RNAs elicit responses with a stronger antiviral component and a lower proinflammatory component. (**A**) Principal component analysis of A549 cells transcriptomes when transfected with scrambled dsRNA (ctrl), RNA-1 (isRNA) or poly(I:C) for 48 hours. N=3. (**B** and **C**) Volcano plots showing significant upregulated genes (red) or downregulated genes (blue) in isRNA transfected (**B**) or poly(I:C) transfected (**C**) A549 cells. Threshold for fold change = 2, threshold for P_adj_ = 0.01. (**D**). Heat map showing top upregulated inflammatory genes and top downregulated genes involved in ion transport and cell-cell adhesion in the poly(I:C) transfected but not in the isRNA transfected A549 cells.

### Broad spectrum inhibition of multiple coronaviruses and influenza A viruses

To explore the potential physiological and clinical relevance of these new RNAs that demonstrated immunostimulatory activities in established cell lines, we investigated whether they can trigger IFN-I responses in human Lung Airway and Alveolus Chip microfluidic culture devices lined by human primary lung bronchial or alveolar epithelium grown under an air-liquid interface in close apposition to a primary pulmonary microvascular endothelium cultured under dynamic fluid flow (Fig. 6A), which have been demonstrated to faithfully recapitulate human organ-level lung physiology and pathophysiology (13,27,28). We observed 12-to 30-fold increases in IFN-β expression compared to a scrambled duplex RNA control when we transfected RNA-1 into human bronchial or alveolar epithelial cells through the air channels of the human Lung Chips (Fig. 6B). In addition, treatment with RNA-1 induced robust (> 40-fold) IFN-β expression in human primary lung endothelium on-chip (Fig. 6B) when it was introduced through the vascular channel.

**Figure 6.**
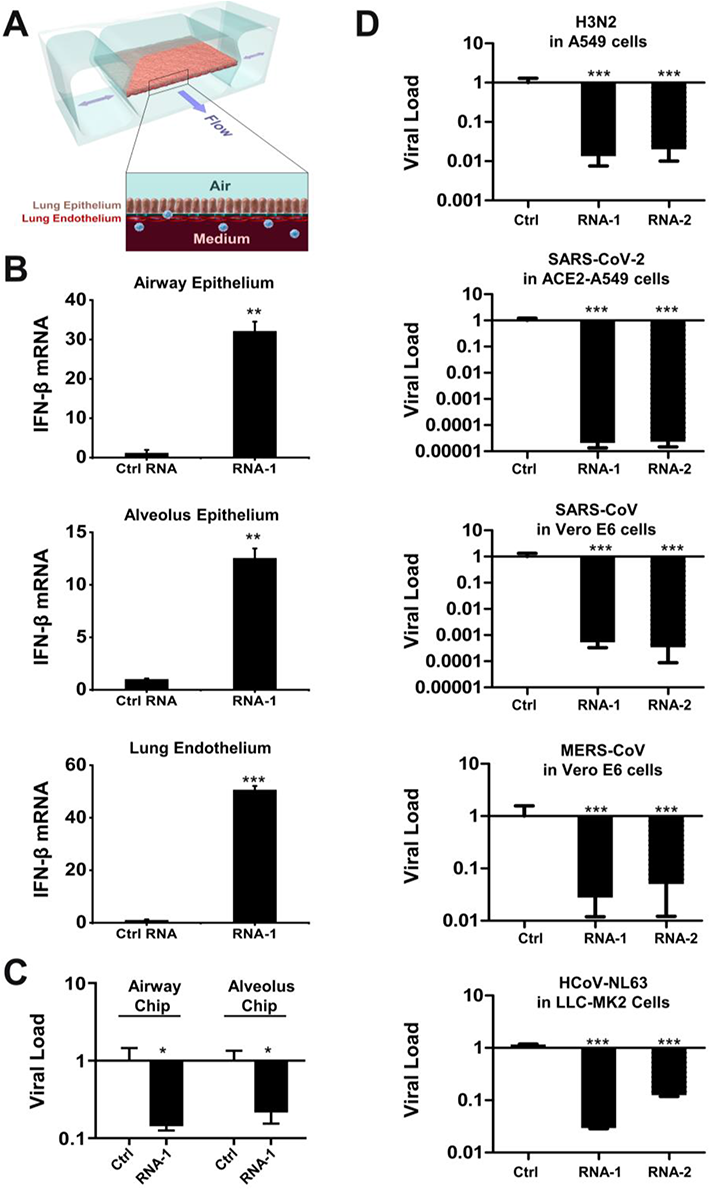
Immunostimulatory RNAs induce IFN-β production in differentiated human lung epithelial and endothelial cells in Organ Chips and exhibit broad spectrum inhibition of infection by H3N2 influenza virus, SARS-CoV-2, SARS-CoV-1, MERS-CoV, and HCoV-NL63. (**A**) Schematic diagram of a cross-section through the human Lung-on-Chip, which faithfully recapitulate human lung physiology and pathophysiology. (**B**) Human Lung Airway and Alveolus Chips were transfected with RNA-1 or scrambled RNA control by perfusion through both channels of the chip and 48 h later, the epithelial and endothelial cells were collected for detection of IFN-β mRNA by qPCR (data are presented as fold change relative to the RNA control; N = 3; *, *p* < 0.05; ***, *p* < 0.001). (**C**) Effects of treatment with RNA-1 or a scrambled control in the human Lung Airway Chips or human Lung Alveolus Chips infected with influenza A/HK/8/68 (H3N2) (MOI = 0.1) at 24 h after RNA-1 treatment. Viral load was determined by quantifying the viral NP gene by qPCR in cell lysates at 48 h after infection. Results are shown as fold change relative to RNA control; N=3; *, *p* < 0.05. (**D**) Treatment with immunostimulatory duplex RNAs resulted in potent inhibition of multiple potential pandemic viruses, including SARS-CoV-2. Indicated cells were treated with RNA-1, RNA-2, or a scrambled control and infected with influenza A/HK/8/68 (H3N2) (MOI = 0.1), SARS-CoV-2 (MOI = 0.05), SARS-CoV-1 (MOI = 0.01), MERS-CoV (MOI = 0.01), and HCoV-NL63 (MOI = 0.002), respectively, at 24 h after RNA transfection. Viral load was determined by quantifying the viral NP gene for H3N2, and the N gene for SARS-CoV-2 and HCoV-NL63 by qPCR in cell lysates at 48 h after infection; viral loads of SARS-CoV and MERS-CoV were determined by plaque assay at 48 h after infection. All results are shown as fold change relative to RNA control; N=3; *, *p* < 0.05; ***, *p* < 0.001.

Given our initial finding that RNA-1 and -2 inhibit infection by H1N1 (Fig. 1A) along with the known antiviral functions of IFN-I/III (29), we next explored the generalizability of these effects. First, we examined the potential of these IFN inducing RNAs to block infection by influenza A/HK/8/68 (H3N2) virus in which cells were transfected with RNAs one day prior to infection, and then with the advent of the COVID-19 pandemic, we extended this work by carrying out similar studies with SARS-CoV-2 and related coronaviruses, SARS-CoV, MERS-CoV, and HCoV-NL63. Analysis with qPCR for viral mRNA revealed that treatment with the immunostimulatory duplex RNAs significantly suppressed infections by H3N2 influenza virus in human Lung Airway and Alveolus Chips (80-90% inhibition) and in A549 cells (>95% inhibition) (Fig. 6C,D), as it did with H1N1 influenza virus in A549 cells (Fig. 1A). Importantly, these same duplex RNAs inhibited MERS-CoV in Vero E6 cells and HCoV-NL63 in LLC-MK2 cells by >90% (Fig. 6D), as well as SARS-CoV in Vero E6 cells by > 1,000-fold (>99.9%) (Fig. 6D). Impressively, they were even more potent inhibitors of SARS-CoV-2 infection, reducing viral load in ACE2 receptor-overexpressing A549 cells by over 10,000-fold (>99.99%) (Fig. 6D and Fig. S9), which is consistent with the observation that SARS-CoV-2 regulates IFN-I/III signaling differently and fails to induce its expression relative to influenza virus and other coronaviruses (30, 31).

## DISCUSSION

In this study, we observed potent stimulation of IFN-I/III signaling by a new class of short duplex RNAs that contain a conserved overhanging sequence motif and terminal monophosphate or hydroxyl groups in a broad spectrum of human cells. Mechanistic exploration revealed that these immunostimulatory RNAs specifically activate the RIG-I/IRF3 pathway by binding directly to RIG-I, even though duplex RNAs with monophosphate groups have been previously shown to antagonize IFN signaling by RNAs with 5’-di or -triphosphates (3, 17). By systematically investigating the effects of various sequences and lengths of these RNAs on IFN-I induction, we identified that the immunostimulatory activity requires a minimal length of 20 bases, in addition to a conserved overhanging sequence motif (sense strand: C, antisense strand: 3’-GGG-5’). This motif mediates the formation of end-to-end duplex RNA dimers via Hoogstein base pairing that enable its binding to RIG-I. In addition, the RNA-mediated IFN-I production that we observed resulted in significant inhibition of infections by multiple human respiratory viruses, including H1N1 and H3N2 influenza viruses, as well as coronaviruses SARS-CoV-2, SARS-CoV-1, MERS-CoV, and HCoV-NL63. Notably, these new immunostimulatory RNAs significantly reduced SARS-CoV-2 viral loads in cell lines and in human Lung Airway and Alveolus Chips containing primary lung epithelial and endothelial cells. These findings raise the possibility that these IFN-inducing immunostimulatory RNAs could offer alternative prophylactic and therapeutic strategies for the current COVID-19 pandemic, in addition to providing potential broad-spectrum protection against a wide range of respiratory viruses that might emerge in the future. In particular, this new duplex RNA approach provides a clear advantage over the commonly used PRR agonist Poly(I:C), as it is fully chemically defined, easier to synthesize, and exerts a more targeted antiviral effect with less proinflammatory activity.

Interestingly, the conserved overhanging motif we identified that contains 5’-C and 3’-GGG ends on the sense and antisense strands, respectively, appears to mediate ‘end-to-end’ dimerization of the duplex RNAs via formation of an intramolecular G-quadruplex generated by the GG overhang as any changes to this motif led to complete loss of immunostimulatory activity. The remaining exposed 5’ ends of the resultant longer dimers, in turn, appears to be responsible for binding directly to RIG-I, which thereby triggers IFN production. Consistent with this hypothesis, N_1_-2’O-methylation at the 5’ end of antisense strand, but not the other ends of the original short dsRNA led to complete loss of the immunostimulatory activity. All of these findings are consistent with previous research demonstrating that RIG-I recognizes the 5’ ends of longer duplex RNAs (3). Notably, similar Hoogsteen-like pairing has been identified between trans U-U base pairs in 5’-UU overhang dsRNA fragments (32); however, our research establishes for the first time that Hoogsteen base pairing can lead to generation of duplex RNAs that are highly effective RIG-I agonists.

siRNA has become a common laboratory tool for gene silencing in biomedical research for almost two decades and a class of drugs that has recently been approved in clinics (11, 12). However, the activation of innate immune responses by siRNAs is challenging their uses in both settings (11,12,33). A number of features that may elicit immune responses by siRNA have been identified (Table S1), for examples, the presence of 5’ triphosphate in siRNA synthesized by phage polymerase (7) or specific sequence motifs in the sense strand of siRNA (8). However, these features do not cover all possible scenarios, including the new immunostimulatory RNAs identified in our study. While optimally designed siRNAs may not have this motif in the overhang because of the potential for the siRNA to be cleaved by RNase at single-stranded G residues (34), our results further highlight the importance to exclude this feature in future siRNA design to alleviate unwanted activation of innate immune responses.

While immune stimulation by siRNAs is undesired in some gene silencing applications, it can be beneficial in others, such as treatment of viral infections or cancer. The IFN response constitutes the major first line of defense against viruses, and these infectious pathogens, including SARS-CoV-2, have evolved various strategies to suppress this response (30, 35). In particular, transcriptomic analyses in both human cultured cells infected with SARS-CoV-2 and COVID-19 patients revealed that SARS-CoV-2 infection produces a unique inflammatory response with very low IFN-I, IFN-III, and associated ISG responses, while still stimulating chemokine and pro-inflammatory cytokine production (30, 35), and this imbalance likely contributes to the increased morbidity and mortality seen in late stage COVID-19 patients. Type I and type III IFN proteins are therefore being evaluated for their efficacy as therapeutics in preclinical models and clinical trials (36–39). Pretreatment with IFN proteins has been shown to reduce viral titers, suggesting that induction of IFN-I responses may represent a potentially effective approach for prophylaxis or early treatment of SARS-CoV-2 infections (40, 41). Triple combination of IFN-β1b, lopinavir-ritonavir, and ribavirin also has been recently reported to shorten the duration of viral shedding and hospital stay in patients with mild to moderate COVID-19 (42).

Consistent with these observations, our results showed that pretreatment with our immunostimulatory RNAs resulted in a dramatic decrease in infection by SARS-CoV-2, as well as SARS-CoV, MERS-CoV, HCoV-NL63 (common cold virus) and H1N1 and H3N2 influenza viruses. Importantly, our immunostimulatory RNAs specifically activate RIG-I/IFN-I pathway but are not recognized by other cellular RNA sensors, such as TLR7, TLR8, MDA5, or TLR3. This is interesting because recent studies show that SARS-CoV-2 inhibits RIG-I signaling and clearance of infection via expression of nsp1 (43). Thus, our results demonstrate that these duplex RNAs can overcome this inhibition, at least in human lung epithelial and endothelial cells maintained in Organ Chips that recapitulate human lung pathophysiology (44, 45).

## MATERIAL AND METHODS

### Cell culture

A549 cells (ATCC CCL-185), A549-Dual^TM^ cells (InvivoGen), RIG-I KO A549-Dual^TM^ cells (InvivoGen), MDA5 KO A549-Dual^TM^ cells (InvivoGen), TLR3 KO A549 cells (Abcam), HEK-Blue^TM^ Null-k cells (InvivoGen, hkb-null1k), HEK-Blue^TM^ hTLR7 cells (InvivoGen, htlr7), THP1-Dual^TM^ cells (InvivoGen, thpd-nifs), THP1-Dual^TM^ KO-TLR8 cells (InvivoGen,kotlr8), MDCK cells (ATCC CRL-2936), and LLC-MK2 cells (ATCC CCL-7.1) were cultured in Dulbecco’s modified Eagle’s medium (DMEM) (Life Technologies) supplemented with 10% fetal bovine serum (FBS) (Life Technologies) and penicillin-streptomycin (Life Technologies). HAP1 cells, IRF3 KO HAP1 cells, and IRF7 KO HAP1 cells were purchased from Horizon Discovery Ltd and cultured in Iscove’s Modified Dulbecco’s Medium (IMDM) (Gibco) supplemented with 10% fetal bovine serum (FBS) (Life Technologies) and penicillin-streptomycin (Life Technologies). All cells were maintained at 37 °C and 5% CO_2_ in a humidified incubator. All cell lines used in this study were free of mycoplasma, as confirmed by the LookOut Mycoplasma PCR Detection Kit (Sigma). Cell lines were authenticated by the ATCC, InvivoGen, Abcam, or Horizon Discovery Ltd. Primary human lung airway epithelial basal stem cells (Lonza, USA) were expanded in 75 cm^2^ tissue culture flasks using airway epithelial cell growth medium (Promocell, Germany) until 60-70% confluent. Primary human alveolar epithelial cells (Cell Biologics, H-6053) were cultured using alveolar epithelial growth medium (Cell Biologics, H6621). Primary human pulmonary microvascular endothelial cells (Lonza, CC-2527, P5) were expanded in 75 cm^2^ tissue culture flasks using human endothelial cell growth medium (Lonza, CC-3202) until 70-80% confluent.

### Viruses

Viruses used in this study include SARS coronavirus-2 (SARS-CoV-2), human coronavirus HCoV-NL63, influenza A/WSN/33 (H1N1), and influenza A/Hong Kong/8/68 (H3N2). SARS-CoV-2 isolate USA-WA1/ 2020 (NR-52281) was deposited by the Center for Disease Control and Prevention, obtained through BEI Resources, NIAID, NIH, and propagated as described previously (30). HCoV-NL63 was obtained from the ATCC and expanded in LLC-MK2 cells. Influenza A/WSN/33 (H1N1) was generated using reverse genetics technique and influenza A/Hong Kong/8/68 (H3N2) was obtained from the ATCC. Both influenza virus strains were expanded in MDCK cells. HCoV-NL63 was titrated in LLC-MK2 cells by Reed-Muench method. Influenza viruses were titrated by plaque formation assay (27). All experiments with native SARS-CoV-2, SARS-CoV, and MERS-CoV were performed in a BSL3 laboratory and approved by our Institutional Biosafety Committee.

### Stimulation of cell lines by transfection

All RNAs and scrambled negative control dsRNA were synthesized by Integrated DNA Technologies, lnc. (IDT). The poly(I:C) was purchased from InvivoGen (Cat# tlrl-picw), which specifically confirmed the absence of contamination by bacterial lipoproteins or endotoxins. Cells were seeded into 6-well plate at 3 × 10^5^ cells/well or 96-well plate at 10^4^ cells/well and cultured for 24 h before transfection. Transfection was performed using TransIT-X2 Dynamic Delivery System (Mirus) according to the manufacturer’s instructions with some modifications. If not indicated otherwise, 6.8 µL of 10 µM RNA stock solution and 5 µL of transfection reagent were added in 200 µL Opti-MEM (Invitrogen) to make the transfection mixture. For transfection in 6-well plate, 200 µL of the transfection mixture was added to each well; for transfection in 96-well plate, 10 µL of the transfection mixture was added to each well. At indicated times after transfection, cell samples were collected and subjected to RNA-seq (Genewiz, lnc.), TMT Mass spectrometry, qRT-PCR, western blot, or Quanti-Luc assay (InvivoGen).

### RNA-seq and Gene ontogeny analysis

RNA-seq was processed by Genewiz using a standard RNA-seq package that includes polyA selection and sequencing on an Illumina HiSeq with 150-bp pair-ended reads. Sequence reads were trimmed to remove possible adapter sequences and nucleotides with poor quality using Trimmomatic v.0.36. The trimmed reads were mapped to the Homo sapiens GRCh38 reference genome using the STAR aligner v.2.5.2b. Unique gene hit counts were calculated by using feature Counts from the Subread package v.1.5.2 followed by differential expression analysis using DESeq2. Gene Ontology analysis was performed using DAVID (46). Volcano plots and heat maps were generated using the EnhancedVolcano R package (47). Raw sequencing data files were deposited on NCBI GEO with the accession number GSE181827.

### Proteomics analysis by Tandem Mass Tag Mass Spectrometry

Cells were harvested on ice. Cells pellets were syringe-lysed in 8 M urea and 200 mM EPPS pH 8.5 with protease inhibitor. BCA assay was performed to determine protein concentration of each sample. Samples were reduced in 5 mM TCEP, alkylated with 10 mM iodoacetamide, and quenched with 15 mM DTT. 100 µg protein was chloroform-methanol precipitated and re-suspended in 100 µL 200 mM EPPS pH 8.5. Protein was digested by Lys-C at a 1:100 protease-to-peptide ratio overnight at room temperature with gentle shaking. Trypsin was used for further digestion for 6 hours at 37°C at the same ratio with Lys-C. After digestion, 30 µL acetonitrile (ACN) was added into each sample to 30% final volume. 200 µg TMT reagent (126, 127N, 127C, 128N, 128C, 129N, 129C, 130N, 130C) in 10 µL ACN was added to each sample. After 1 hour of labeling, 2 µL of each sample was combined, desalted, and analyzed using mass spectrometry. Total intensities were determined in each channel to calculate normalization factors. After quenching using 0.3% hydroxylamine, eleven samples were combined in 1:1 ratio of peptides based on normalization factors. The mixture was desalted by solid-phase extraction and fractionated with basic pH reversed phase (BPRP) high performance liquid chromatography (HPLC), collected onto a 96 six well plate and combined for 24 fractions in total. Twelve fractions were desalted and analyzed by liquid chromatography-tandem mass spectrometry (LC-MS/MS) (48).

Mass spectrometric data were collected on an Orbitrap Fusion Lumos mass spectrometer coupled to a Proxeon NanoLC-1200 UHPLC. The 100 µm capillary column was packed with 35 cm of Accucore 50 resin (2.6 μm, 150Å; ThermoFisher Scientific). The scan sequence began with an MS1 spectrum (Orbitrap analysis, resolution 120,000, 375−1500 Th, automatic gain control (AGC) target 4E5, maximum injection time 50 ms). SPS-MS3 analysis was used to reduce ion interference (49, 50). The top ten precursors were then selected for MS2/MS3 analysis. MS2 analysis consisted of collision-induced dissociation (CID), quadrupole ion trap analysis, automatic gain control (AGC) 2E4, NCE (normalized collision energy) 35, q-value 0.25, maximum injection time 35ms), and isolation window at 0.7. Following acquisition of each MS2 spectrum, we collected an MS3 spectrum in which multiple MS2 fragment ions are captured in the MS3 precursor population using isolation waveforms with multiple frequency notches. MS3 precursors were fragmented by HCD and analyzed using the Orbitrap (NCE 65, AGC 1.5E5, maximum injection time 120 ms, resolution was 50,000 at 400 Th).

Mass spectra were processed using a Sequest-based pipeline (51). Spectra were converted to mzXML using a modified version of ReAdW.exe. Database searching included all entries from the Human UniProt database (downloaded: 2014-02-04) This database was concatenated with one composed of all protein sequences in the reversed order. Searches were performed using a 50 ppm precursor ion tolerance for total protein level analysis. The product ion tolerance was set to 0.9 Da. TMT tags on lysine residues and peptide N termini (+229.163 Da) and carbamidomethylation of cysteine residues (+57.021 Da) were set as static modifications, while oxidation of methionine residues (+15.995 Da) was set as a variable modification.

Peptide-spectrum matches (PSMs) were adjusted to a 1% false discovery rate (FDR) (52, 53). PSM filtering was performed using a linear discriminant analysis (LDA), as described previously (51), while considering the following parameters: XCorr, ΔCn, missed cleavages, peptide length, charge state, and precursor mass accuracy. For TMT-based reporter ion quantitation, we extracted the summed signal-to-noise (S:N) ratio for each TMT channel and found the closest matching centroid to the expected mass of the TMT reporter ion. For protein-level comparisons, PSMs were identified, quantified, and collapsed to a 1% peptide false discovery rate (FDR) and then collapsed further to a final protein-level FDR of 1%, which resulted in a final peptide level FDR of < 0.1%. Moreover, protein assembly was guided by principles of parsimony to produce the smallest set of proteins necessary to account for all observed peptides. Proteins were quantified by summing reporter ion counts across all matching PSMs, as described previously (51). PSMs with poor quality, MS3 spectra with TMT reporter summed signal-to-noise of less than 100, or having no MS3 spectra were excluded from quantification (54). Each reporter ion channel was summed across all quantified proteins and normalized assuming equal protein loading of all tested samples. Raw data were submitted to ProteomeXchange via the PRIDE database with the accession PXD027838.

### qRT-PCR

Total RNA was extracted from cells using RNeasy Plus Mini Kit (QiaGen, Cat#74134) according to the manufacturer’s instructions. cDNA was then synthesized using AMV reverse transcriptase kit (Promega) according to the manufacturer’s instructions. To detect gene levels, quantitative real-time PCR was carried out using the GoTaq qPCR Master Mix kit (Promega) with 20 µL of reaction mixture containing gene-specific primers or the PrimePCR assay kit (Bio-Rad) according the manufacturers’ instructions. The expression levels of target genes were normalized to GAPDH.

### Antibodies and Western blotting

The antibodies used in this study were anti-IRF3 (Abcam, ab68481), anti-IRF3 (Phospho S396) (Abcam, ab138449), anti-GAPDH (Abcam, ab9385), and Goat anti-Rabbit IgG H&L (HRP) (Abcam, ab205718). Cells were harvested and lysed in RIPA buffer (Thermo Scientific, Cat#89900) supplemented with Halt^TM^ protease and phosphatase inhibitor cocktail (Thermo Scientific, Cat#78440) on ice. The cell lysates were subject to western blotting. GAPDH was used as a loading control.

### Confocal immunofluorescence microscopy

Cells were rinsed with PBS, fixed with 4% paraformaldehyde (Alfa Aesar) for 30 min, permeabilized with 0.1% Triton X-100 (Sigma-Aldrich) in PBS (PBST) for 10 min, blocked with 10% goat serum (Life Technologies) in PBST for 1 h at room temperature, and incubated with anti-IRF3 (Phospho S396) (Abcam, ab138449) antibody diluted in blocking buffer (1% goat serum in PBST) overnight at 4 °C, followed by incubation with Alexa Fluor 488 conjugated secondary antibody (Life Technologies) for 1 h at room temperature; nuclei were stained with DAPI (Invitrogen) after secondary antibody staining. Fluorescence imaging was carried out using a confocal laser-scanning microscope (SP5 X MP DMI-6000, Germany) and image processing was done using Imaris software (Bitplane, Switzerland).

### Surface plasmon resonance

The interactions between duplex RNA-1 and cellular RNA sensor molecules (RIG-I (Abcam, Cat# ab271486), MDA5 (Creative-Biomart, Cat# IFIH1-1252H), and TLR3 (Abcam, Cat# ab73825)) were analyzed by SPR with the Biacore T200 system (GE Healthcare) at 25 °C (Creative-Biolabs Inc.). RNA-1 conjugated with biotin (synthesized by IDT lnc.) was immobilized on a SPR sensor chip, with final levels of ∼60 response units (RU). Various concentrations of the RNA sensors diluted in running buffer (10 × HBS-EP+; GE Healthcare, Cat# BR100669) were injected as analytes at a flow rate of 30 μl/min, a contact time of 180 s, and a dissociation time of 300 s. The surface was regenerated with 2 M NaCl for 60 s. Data analysis was performed on the Biacore T200 computer with the Biacore T200 evaluation software.

### Organ Chip Culture

Microfluidic two-channel Organ Chip devices and automated ZOE® instruments used to culture them were obtained from Emulate Inc (Boston, MA, USA). Our methods for culturing human Lung Airway Chips (27, 28) and Lung Alveolus Chips have been described previously. In this study, we slightly modified the Alveolus Chip method by coating the inner channels of the devices with 200 µg/ml Collagen IV (5022-5MG, Advanced Biomatrix) and 15 μg/ml of laminin (L4544-100UL, Sigma) at 37°C overnight, and the next day (day 1) sequentially seeding primary human lung microvascular endothelial cells (Lonza, CC-2527, P5) and primary human lung alveolar epithelial cells (Cell Biologics, H-6053) in the bottom and top channels of the chip at a density of 8 and 1.6 x 10^6^ cells/ml, respectively, under static conditions. On day 2, the chips were inserted into Pods® (Emulate Inc.), placed within the ZOE® instrument, and the apical and basal channels were respectively perfused (60 μL/hr) with epithelial growth medium (Cell Biologics, H6621) and endothelial growth medium (Lonza, CC-3202). On day 5, 1 uM dexamethasone was added to the apical medium to enhance barrier function. On day 7, an air-liquid interface (ALI) was introduced into the epithelial channel by removing all medium from this channel while continuing to feed all cells through the medium perfused through the lower vascular channel, and this medium was changed to EGM-2MV with 0.5% FBS on day 9. Two days later, the ZOE® instrument was used to apply cyclic (0.25 Hz) 5% mechanical strain to the engineered alveolar-capillary interface to mimic lung breathing on-chip. RNAs were transfected on Day 15.

### RNA transfection in human Lung Airway and Alveolus Chips

Human Airway or Alveolus Chips were transfected with duplex RNAs by adding the RNA and transfection reagent (Lipofectamine RNAiMAX) mixture into the apical and basal channels of the Organ Chips and incubating for 6 h at 37°C under static conditions before reestablishing an ALI. Tissues cultured on-chip were collected by RNeasy Micro Kit (QiaGen) at 48 h post-transfection by first introducing 100 ul lysis buffer into the apical channel to lyse epithelial cells and then 100 ul into the basal channel to lyse endothelial cells. Lysates were subjected to qPCR analysis of IFN-β gene expression.

### Native SARS-CoV-2 infection and inhibition by RNA treatment

ACE2-expressing A549 cells (a gift from Brad Rosenberg) were transfected with indicated RNAs. 24 h post-transfection, the transfected ACE2-A549 cells were infected with SARS-CoV-2 (MOI = 0.05) for 48 hours. Cells were harvested in Trizol (Invitrogen) and total RNA was isolated and DNAse-I treated using Zymo RNA Miniprep Kit according to the manufacturer’s protocol. qRT-PCR for α-tubulin (Forward: 5’-GCCTGGACCACAAGTTTGAC-3’; Reverse: 3’-TGAAATTCTGGGAGCATGAC-5’) and SARS-CoV-2 N mRNA (Forward: 5’-CTCTTGTAGATCTGTTCTCTAAACGAAC-3’; Reverse: 3’-GGTCCACCAAACGTAATGCG-5’) were performed using KAPA SYBR FAST ONE-STEP qRT-PCR kits (Roche) according to manufacturer’s instructions on a Lightcycler 480 Instrument-II (Roche).

### Native SARS-CoV-1 and MERS-CoV infection and inhibition by RNA treatment

Vero E6 cells (ATCC# CRL 1586) were cultured in DMEM (Quality Biological), supplemented with 10% (v/v) fetal bovine serum (Sigma), 1% (v/v) penicillin/streptomycin (Gemini Bio-products) and 1% (v/v) L-glutamine (2 mM final concentration, Gibco). Cells were maintained at 37°C (5% CO_2_). Vero E6 cells were plated at 1.5x 10^5^ cells per well in a six well plate two days prior to transfection. The RNA-1, RNA-2, and scrambled control RNA were transfected into each well using the Transit X2 delivery system (MIRUS; MIR6003) in OptiMEM (Gibco 31985-070). SARS-CoV (Urbani strain, BEI#NR-18925) and MERS-CoV (Jordan strain, provided by NIH) were added at MOI 0.01. At 72 hours post infection, medium was collected and used for a plaque assay to quantify PFU/mL of virus.

### Quantification and Statistical Analysis

All data are expressed as mean ± standard deviation (SD). N represents biological replicates. Statistical significance of differences in the *in vitro* experiments was determined by employing the paired two-tailed Student t-test when comparing the difference between two groups and one-way ANOVA with multiple comparison when comparing the samples among groups with more than two samples. For all experiments, differences were considered statistically significant for *p* < 0.05 (*, *p* < 0.05; **, *p* < 0.01; ***, *p* < 0.001; n.s., not significant).

## Acknowledgements

We thank IDT, lnc. for synthesizing the RNA oligonucleotides, X. Song and H. Queen at Creative-Biolabs Inc. for carrying SPR experiments and Genewiz, lnc. for carrying out the RNA-seq work. This work was supported by the National Institute of Health (NCATS UH3-HL-141797 to D.E.I.) and the Defense Advanced Research Projects Agency under Cooperative Agreements (HR00111920008 and HR0011-20-2-0040 to D.E.I.).

## Availability

Sharing of materials will be subject to standard material transfer agreements. The raw source data of RNA-seq and TMT Mass Spectrometry have been deposited in Gene Expression Omnibus database under the accession code GSE181827 and PXD027838. Additional data are presented in the Supplementary Materials.

## Conflict of Interest

D.E.I. is a founder, board member, SAB chair, and equity holder in Emulate Inc. D.E.I., L. S., H. B., C.O., and R.P. are inventors on relevant patent applications held by Harvard University.

## Author contributions

L.S., H.B., and D.E.I. conceived this study. L.S. and H.B. conducted in vitro experiments and analyzed data with assistance from C.O., A.J., C.B., W.C., and R.K.P.; T.Z. and S.P.G. performed TMT Mass Spectrometry and data analysis. F.H. performed the native gel electrophoresis experiments. Y.Y. performed the analysis of CCC sequence distribution in human mRNAs and lncRNAs. T.J., J.L., M.F., and B.R.T. performed the experiments of SARS-CoV-2, SARS-CoV, MERS-CoV viruses. A.N. performed western blotting experiments. R.P. assisted in the propagation and characterization of HCoV-NL63 virus. X. Song and H. Queen at Creative-Biolabs Inc. carried SPR experiments. L.S., H.B., and D.E.I. wrote the manuscript with input from other authors.

## Supplementary Data

**Figure S1.**
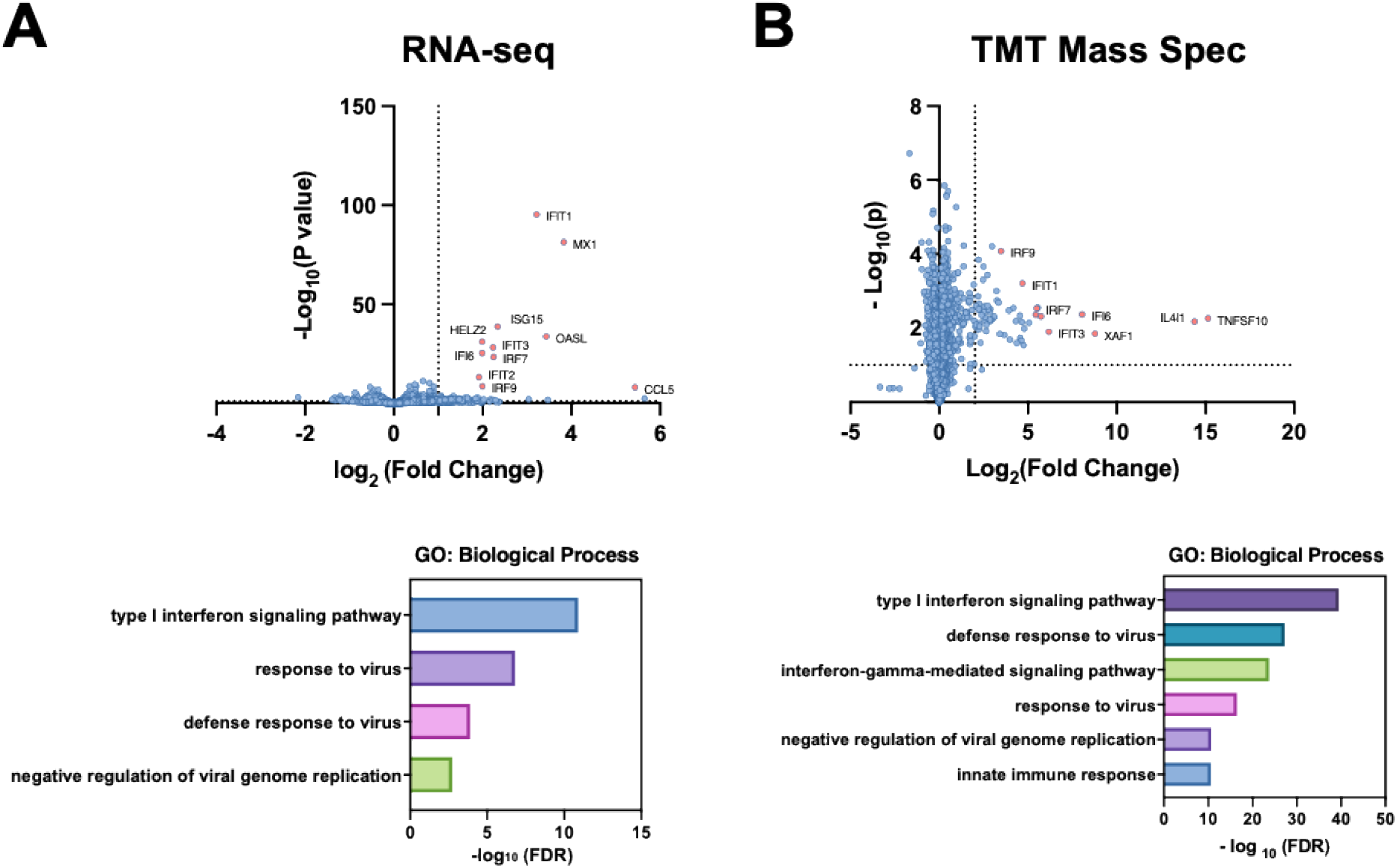
Profiling the effects of RNA-2 by RNA-seq and TMT mass spectrometry. A549 cells were transfected with RNA-2 or scrambled RNA control, cell lysates were collected at 48 h, and analyzed by RNA-seq (left) or TMT Mass Spec (right). Differentially expressed genes (DEGs) or proteins are shown in volcano plots (top) and GO Enrichment analysis was performed for the DEGs (bottom) (N = 3). Plot (top) and GO Enrichment analysis was performed for the differentially expressed proteins (bottom) (N = 3).

**Figure S2.**
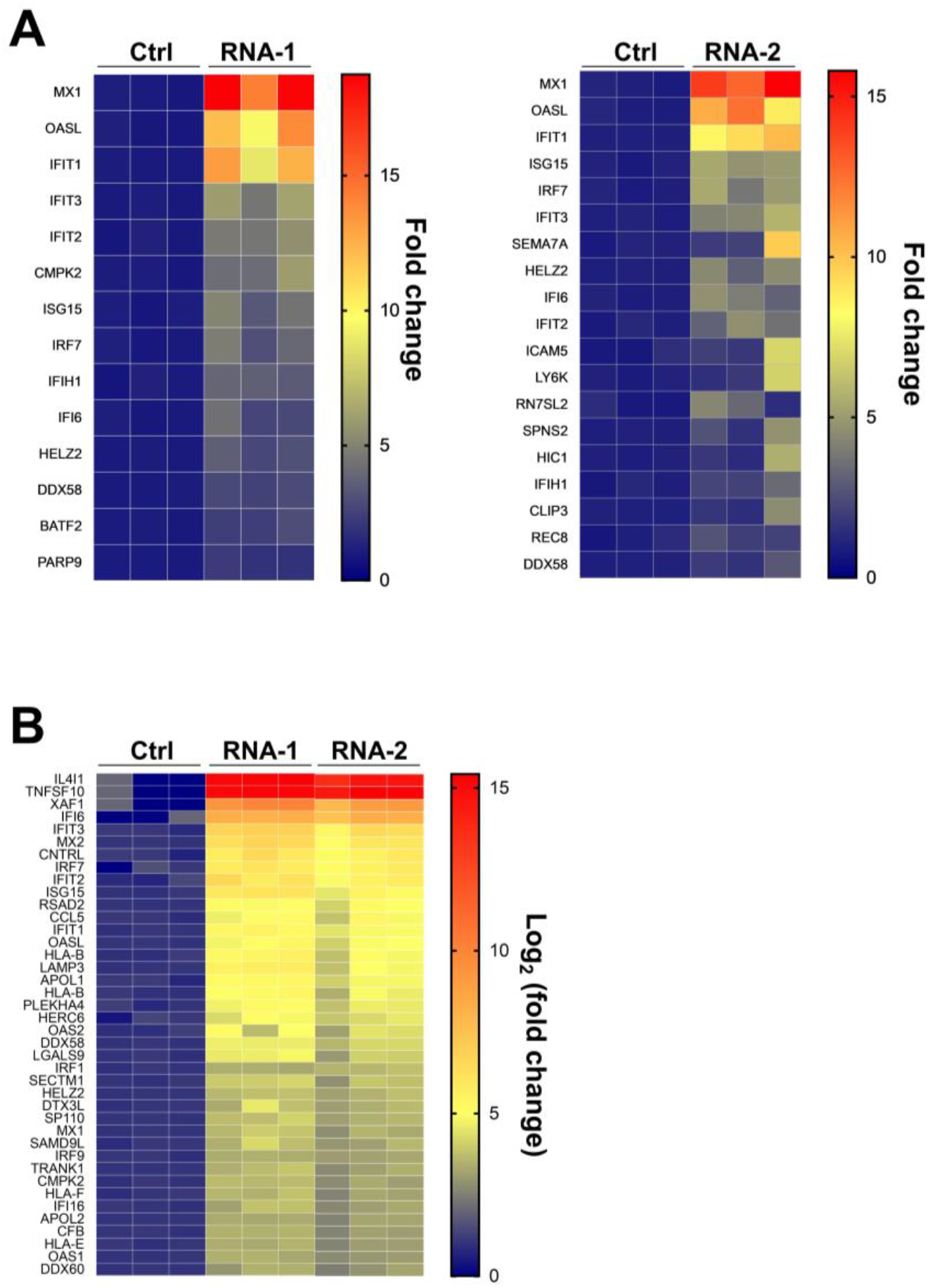
Heat maps showing the effects of immunostimulatory RNAs on IFN pathway-relevant gene levels. DEGs from RNA-seq (**A**) and differentially expressed proteins from TMT Mass Spec analyses (**B**) shown in Fig. 1B and fig. S1 are presented here as heat maps (gene levels of the scrambled RNA control were set as 1; N = 3).

**Figure S3.**
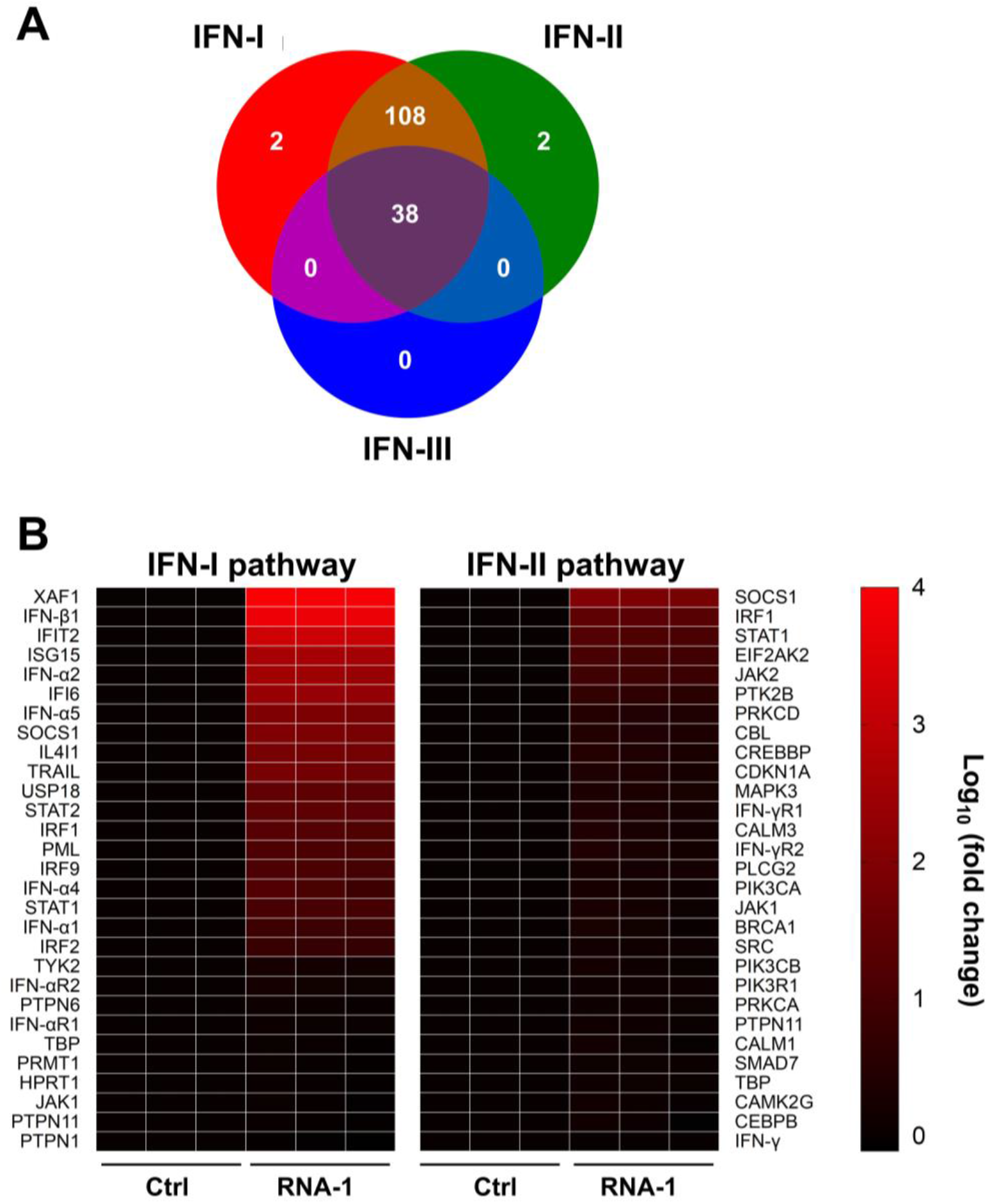
RNA-induced gene expression associated with type I interferon pathway. (**A**) Venn diagram showing differentially expressed ISGs from TMT Mass Spec by RNA-1 belong to type I or type II interferon stimulated genes. (**B**) Heat map of qPCR results showing RNA-I preferentially activates type I interferon pathway. A549 cells were transfected with RNA-1 or scrambled dsRNA control, collected at 48 hr and analyzed by qPCR (expression levels were normalized to GAPDH; gene levels induced by the RNA control were set as 1; N = 3).

**Figure S4.**
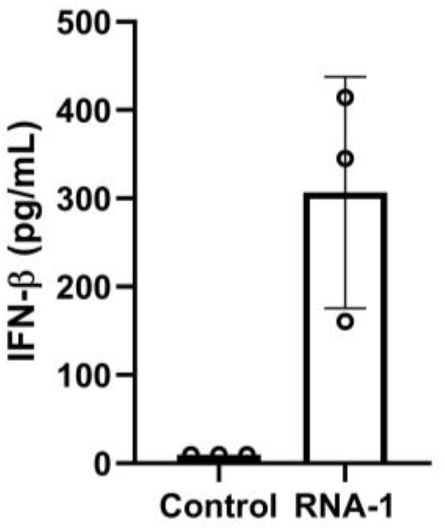
The levels of IFN-β protein induced by RNA-1. A549 cells were transfected with RNA-1 (34 nM) for 48 h, and then supernatants were collected for detection of IFN-β using ELISA. Scrambled RNA control NC-1 is used as negative (N = 3).

**Figure S5.**
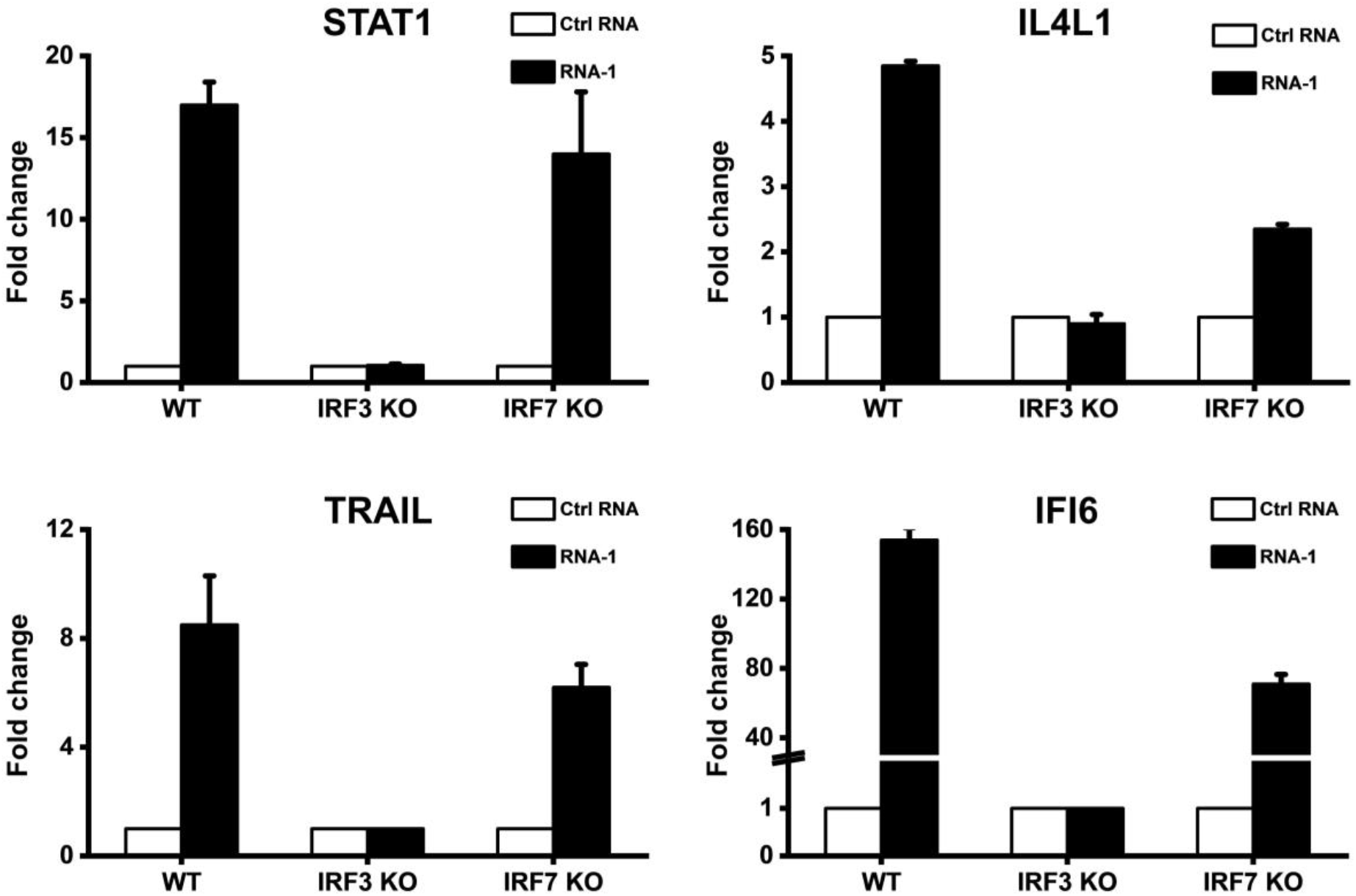
IRF3 knockout abolished the ability of immunostimulatory RNAs to induce IFN-I pathway associated genes. Wild-type (WT) HAP1 cells, IRF3 knockout HAP1 cells, or IRF7 knockout HAP1 cells were transfected with RNA-1 or a scrambled RNA control and STAT1, IL4L1, TRAIL, and IFI6 mRNA levels were quantified by qPCR at 48 h post transfection. Data are presented as fold change relative to RNA control (N = 3).

**Figure S6.**
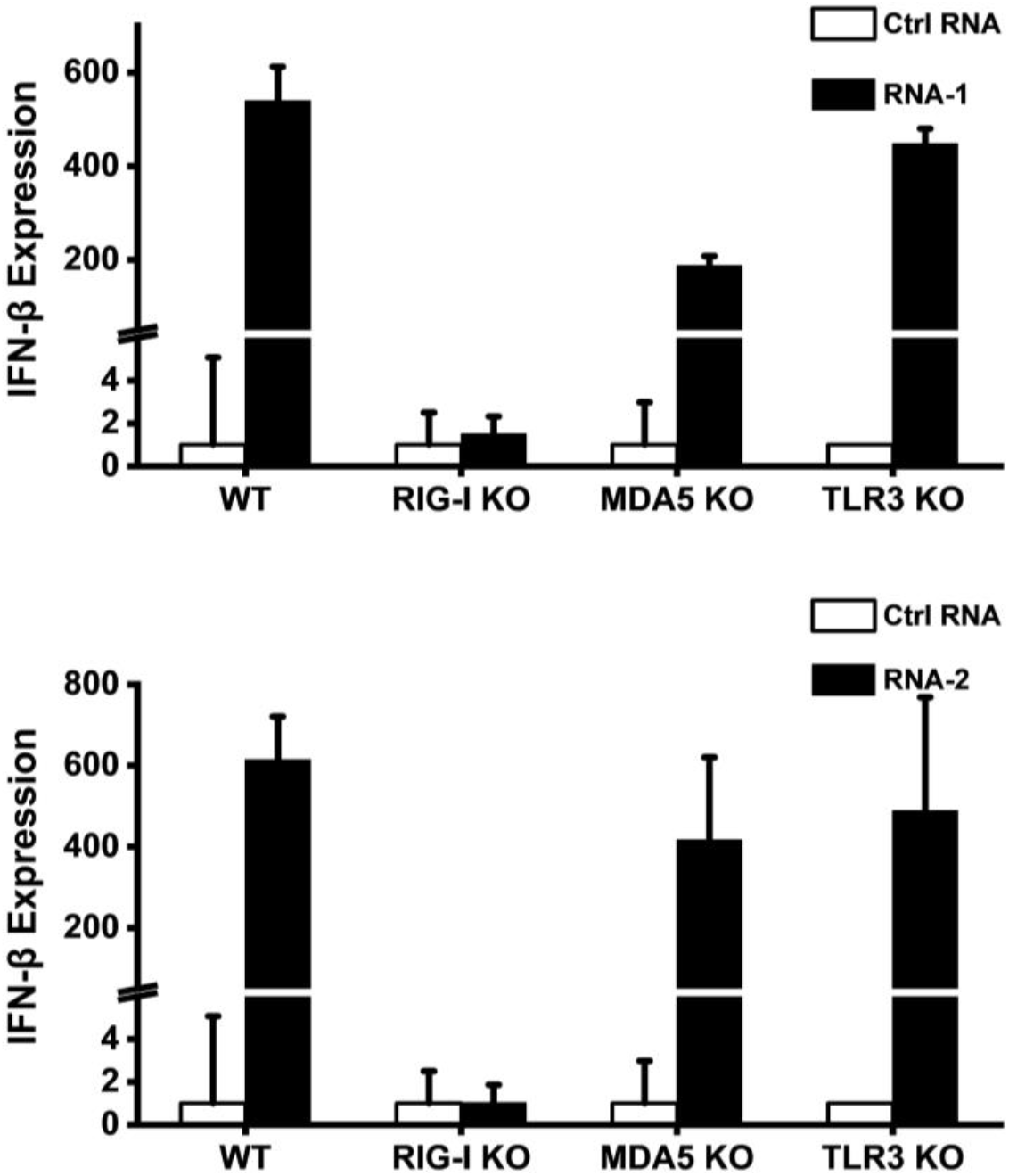
RIG-I knockout abolished the induction effects of the immunostimulatory RNAs on IFN-β. Wild-type (WT) A549-Dual cells, RIG-I knockout A549-Dual cells, MDA5 knockout A549-Dual cells, or TLR3 knockout A549 cells were transfected with RNA-1, RNA-2, or a scramble RNA control and IFN-β mRNA levels were detected by Quanti-Luc assay in WT, RIG-I KO, and MDA5 KO A549-Dual cells or qPCR in TLR3 KO A549 cells at 48 h post transfection. Data are shown as fold change relative to the scrambled RNA control (N = 6).

**Figure S7.**
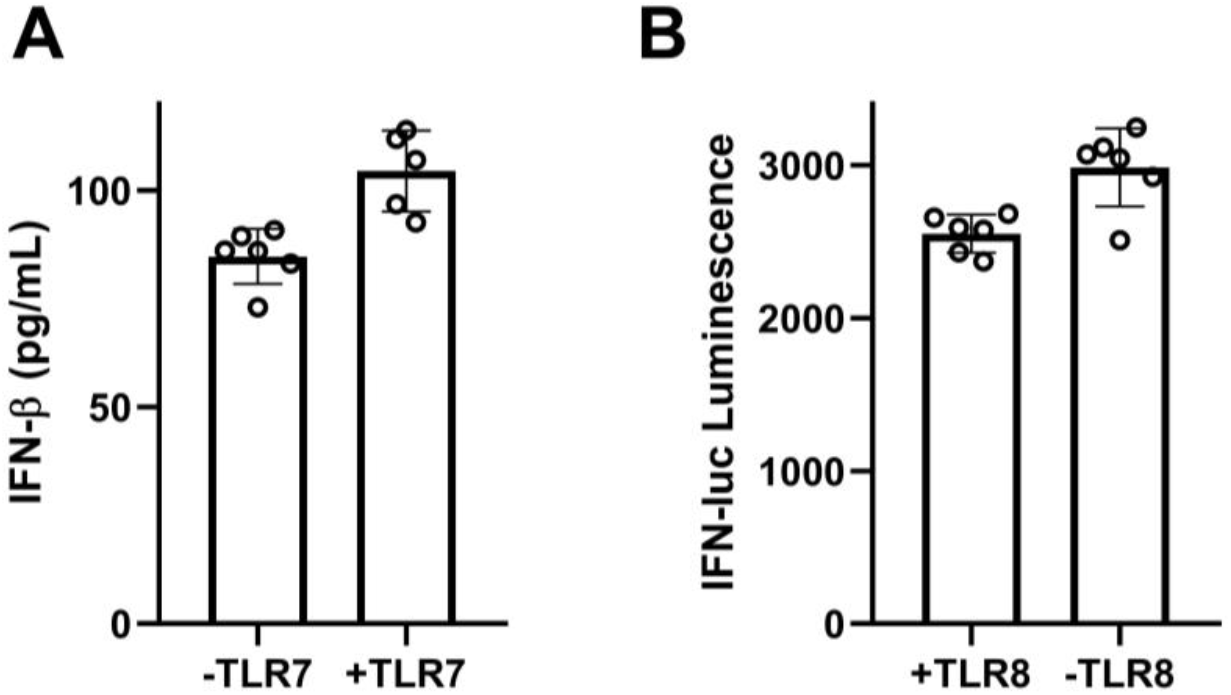
TLR7/8 knockout or overexpression did not have effect on the immunostimulatory activity of RNA-1. (**A**) Graph showing that the overexpression of TLR7 in HEK cells had no effect on production of IFN-β induced by RNA-1. (**B**) Graph showing that the knockout of TLR8 in THP1 cells had no effect on IFN production induced by RNA1. These cell lines are commercial and could be purchased from InvivoGen.

**Figure S8.**
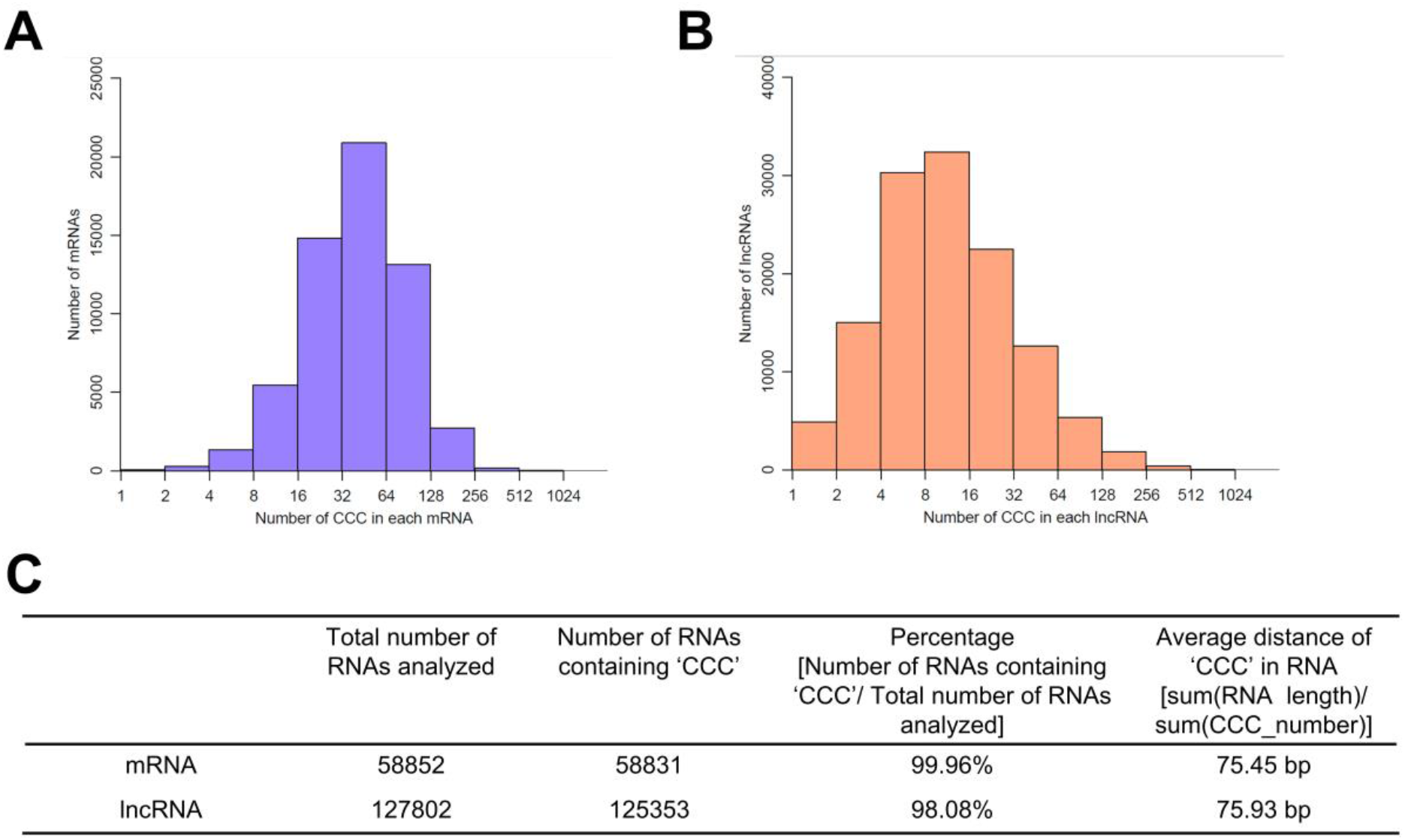
‘CCC’ motif is widely distributed in human genome. (**A**) Graph showing the distribution of the number of CCC sequences in human mRNAs (retrieved from UCSC hg38 refGene with prefix NM). (**B**) Graph showing the distribution of the number of CCC sequences in human lncRNAs (retrieved from lncipedia). (**C**) Table showing the percentage of human mRNAs and lncRNAs containing the CCC motif and their average density.

**Figure S9.**
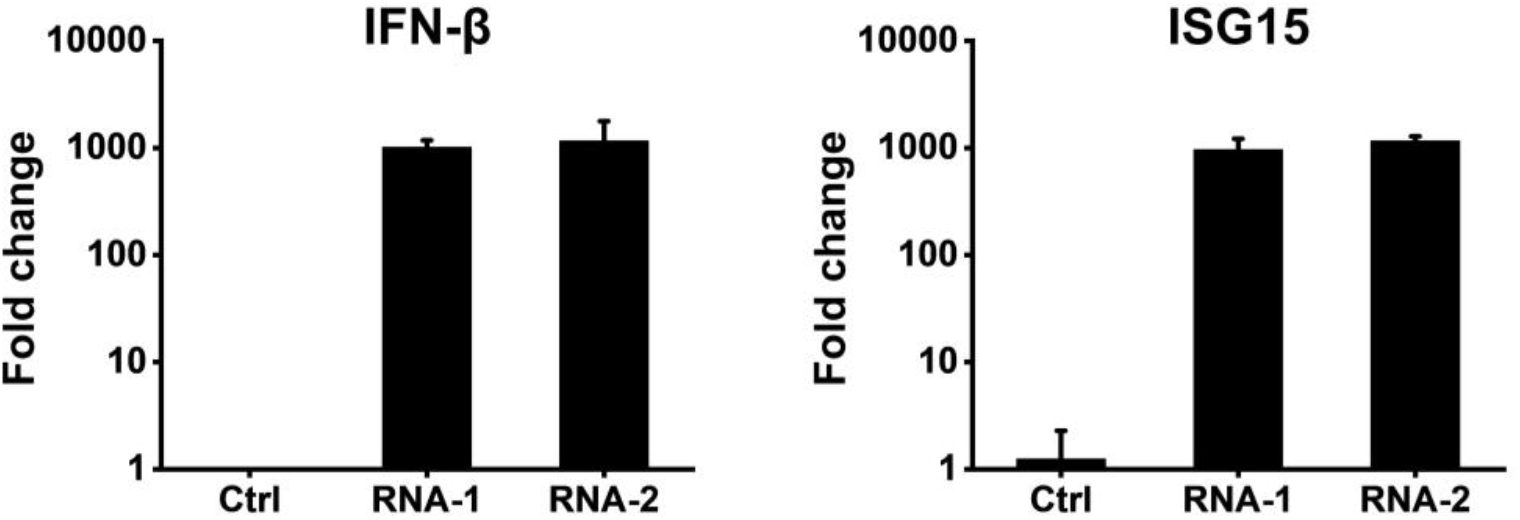
Immunostimulatory RNA-mediated production of IFN in ACE2-overexpressing A549 cells. IFN-β and ISG15 levels were detected in cells transfected with RNA-1, RNA-2, or scramble dsRNA control by qPCR at 48 h post-transfection. The IFN-β or ISG15 level induced by the scramble dsRNA control was set as 1. Data are shown as fold change relative to the control (N = 3).

**Table S1.**
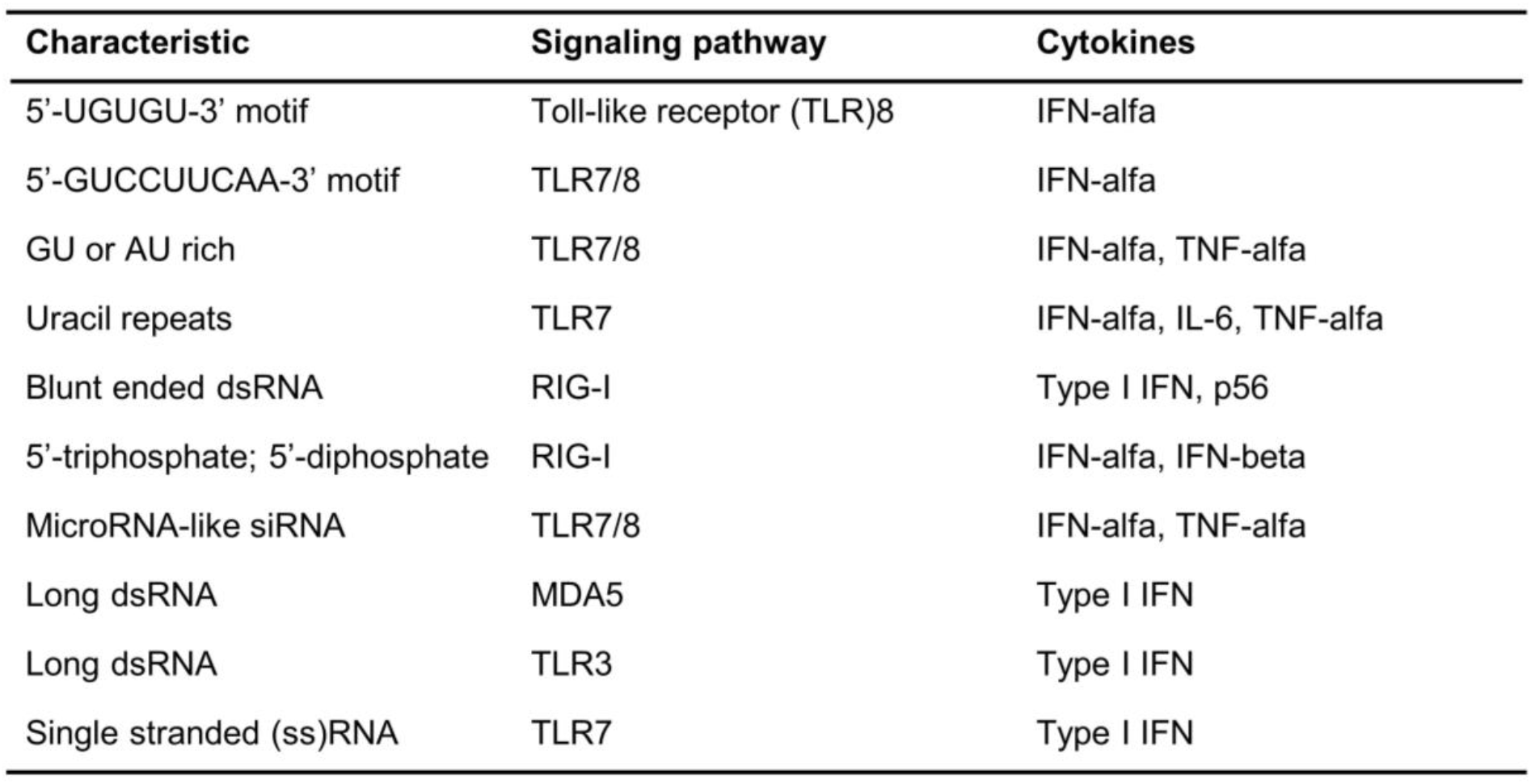
Summary of characteristics of reported immunostimulatory RNAs.

## REFERENCES

1. Schlee, M. and Hartmann, G. (2016) Discriminating self from non-self in nucleic acid sensing. Nat Rev Immunol, 16, 566–580.

2. Kato, H., Takeuchi, O., Mikamo-Satoh, E., Hirai, R., Kawai, T., Matsushita, K., Hiiragi, A., Dermody, T.S., Fujita, T. and Akira, S. (2008) Length-dependent recognition of double-stranded ribonucleic acids by retinoic acid-inducible gene-I and melanoma differentiation-associated gene 5. J Exp Med, 205, 1601–1610.

3. Ren, X., Linehan, M.M., Iwasaki, A. and Pyle, A.M. (2019) RIG-I Selectively Discriminates against 5’-Monophosphate RNA. Cell Rep, 26, 2019–2027 e2014.

4. Ren, X., Linehan, M.M., Iwasaki, A. and Pyle, A.M. (2019) RIG-I Recognition of RNA Targets: The Influence of Terminal Base Pair Sequence and Overhangs on Affinity and Signaling. Cell Rep, 29, 3807–3815 e3803.

5. Jiang, F., Ramanathan, A., Miller, M.T., Tang, G.Q., Gale, M., Jr., Patel, S.S. and Marcotrigiano, J. (2011) Structural basis of RNA recognition and activation by innate immune receptor RIG-I. Nature, 479, 423–427.

6. Marques, J.T. and Williams, B.R. (2005) Activation of the mammalian immune system by siRNAs. Nat Biotechnol, 23, 1399–1405.

7. Kim, D.H., Longo, M., Han, Y., Lundberg, P., Cantin, E. and Rossi, J.J. (2004) Interferon induction by siRNAs and ssRNAs synthesized by phage polymerase. Nat Biotechnol, 22, 321–325.

8. Hornung, V., Guenthner-Biller, M., Bourquin, C., Ablasser, A., Schlee, M., Uematsu, S., Noronha, A., Manoharan, M., Akira, S., de Fougerolles, A. et al. (2005) Sequence-specific potent induction of IFN-alpha by short interfering RNA in plasmacytoid dendritic cells through TLR7. Nat Med, 11, 263–270.

9. Sledz, C.A., Holko, M., de Veer, M.J., Silverman, R.H. and Williams, B.R. (2003) Activation of the interferon system by short-interfering RNAs. Nat Cell Biol, 5, 834–839.

10. Robbins, M., Judge, A., Ambegia, E., Choi, C., Yaworski, E., Palmer, L., McClintock, K. and MacLachlan, I. (2008) Misinterpreting the therapeutic effects of small interfering RNA caused by immune stimulation. Hum Gene Ther, 19, 991–999.

11. Setten, R.L., Rossi, J.J. and Han, S.P. (2019) The current state and future directions of RNAi-based therapeutics. Nat Rev Drug Discov, 18, 421–446.

12. Meng, Z. and Lu, M. (2017) RNA Interference-Induced Innate Immunity, Off-Target Effect, or Immune Adjuvant? Front Immunol, 8, 331.

13. Benam, K.H., Villenave, R., Lucchesi, C., Varone, A., Hubeau, C., Lee, H.H., Alves, S.E., Salmon, M., Ferrante, T.C., Weaver, J.C. et al. (2016) Small airway-on-a-chip enables analysis of human lung inflammation and drug responses in vitro. Nat Methods, 13, 151–157.

14. Huh, D., Matthews, B.D., Mammoto, A., Montoya-Zavala, M., Hsin, H.Y. and Ingber, D.E. (2010) Reconstituting organ-level lung functions on a chip. Science, 328, 1662–1668.

15. Si, L., Bai, H., Rodas, M., Cao, W., Oh, C.Y., Jiang, A., Moller, R., Hoagland, D., Oishi, K., Horiuchi, S. et al. (2021) A human-airway-on-a-chip for the rapid identification of candidate antiviral therapeutics and prophylactics. Nat Biomed Eng.

16. Kim, D.H., Behlke, M.A., Rose, S.D., Chang, M.S., Choi, S. and Rossi, J.J. (2005) Synthetic dsRNA Dicer substrates enhance RNAi potency and efficacy. Nat Biotechnol, 23, 222–226.

17. Goubau, D., Schlee, M., Deddouche, S., Pruijssers, A.J., Zillinger, T., Goldeck, M., Schuberth, C., Van der Veen, A.G., Fujimura, T., Rehwinkel, J. et al. (2014) Antiviral immunity via RIG-I-mediated recognition of RNA bearing 5’-diphosphates. Nature, 514, 372–375.

18. Tissari, J., Siren, J., Meri, S., Julkunen, I. and Matikainen, S. (2005) IFN-alpha enhances TLR3-mediated antiviral cytokine expression in human endothelial and epithelial cells by up-regulating TLR3 expression. Journal of immunology, 174, 4289–4294.

19. Liu, S., Cai, X., Wu, J., Cong, Q., Chen, X., Li, T., Du, F., Ren, J., Wu, Y.T., Grishin, N.V. et al. (2015) Phosphorylation of innate immune adaptor proteins MAVS, STING, and TRIF induces IRF3 activation. Science, 347, aaa2630.

20. Wang, P., Xu, J., Wang, Y. and Cao, X. (2017) An interferon-independent lncRNA promotes viral replication by modulating cellular metabolism. Science, 358, 1051–1055.

21. Zhou, Y., Li, M., Xue, Y., Li, Z., Wen, W., Liu, X., Ma, Y., Zhang, L., Shen, Z. and Cao, X. (2019) Interferon-inducible cytoplasmic lncLrrc55-AS promotes antiviral innate responses by strengthening IRF3 phosphorylation. Cell Res, 29, 641–654.

22. Fitzgerald, K.A., McWhirter, S.M., Faia, K.L., Rowe, D.C., Latz, E., Golenbock, D.T., Coyle, A.J., Liao, S.M. and Maniatis, T. (2003) IKKepsilon and TBK1 are essential components of the IRF3 signaling pathway. Nat Immunol, 4, 491–496.

23. Chow, K.T., Gale, M., Jr. and Loo, Y.M. (2018) RIG-I and Other RNA Sensors in Antiviral Immunity. Annu Rev Immunol, 36, 667–694.

24. Schuberth-Wagner, C., Ludwig, J., Bruder, A.K., Herzner, A.M., Zillinger, T., Goldeck, M., Schmidt, T., Schmid-Burgk, J.L., Kerber, R., Wolter, S. et al. (2015) A Conserved Histidine in the RNA Sensor RIG-I Controls Immune Tolerance to N1-2’O-Methylated Self RNA. Immunity, 43, 41–51.

25. Lyu, K., Chow, E.Y., Mou, X., Chan, T.F. and Kwok, C.K. (2021) RNA G-quadruplexes (rG4s): genomics and biological functions. Nucleic Acids Res, 49, 5426–5450.

26. Blanco-Melo, D., Nilsson-Payant, B.E., Liu, W.C., Uhl, S., Hoagland, D., Moller, R., Jordan, T.X., Oishi, K., Panis, M., Sachs, D. et al. (2020) Imbalanced Host Response to SARS-CoV-2 Drives Development of COVID-19. Cell.

27. Si, L., Bai, H., Rodas, M., Cao, W., Oh, C.Y., Jiang, A., Nurani, A., Zhu, D.Y., Goyal, G., Gilpin, S.E. et al. (2020) Human organs-on-chips as tools for repurposing approved drugs as potential influenza and COVID19 therapeutics in viral pandemics. bioRxiv.

28. Si, L., Prantil-Baun, R., Benam, K.H., Bai, H., Rodas, M., Burt, M. and Ingber, D.E. (2019) Discovery of influenza drug resistance mutations and host therapeutic targets using a human airway chip. bioRxiv.

29. Mesev, E.V., LeDesma, R.A. and Ploss, A. (2019) Decoding type I and III interferon signalling during viral infection. Nat Microbiol, 4, 914–924.

30. Blanco-Melo, D., Nilsson-Payant, B.E., Liu, W.C., Uhl, S., Hoagland, D., Moller, R., Jordan, T.X., Oishi, K., Panis, M., Sachs, D. et al. (2020) Imbalanced Host Response to SARS-CoV-2 Drives Development of COVID-19. Cell, 181, 1036–1045 e1039.

31. Galani, I.E., Rovina, N., Lampropoulou, V., Triantafyllia, V., Manioudaki, M., Pavlos, E., Koukaki, E., Fragkou, P.C., Panou, V., Rapti, V. et al. (2021) Untuned antiviral immunity in COVID-19 revealed by temporal type I/III interferon patterns and flu comparison. Nat Immunol, 22, 32–40.

32. Wahl, M.C., Rao, S.T. and Sundaralingam, M. (1996) The structure of r(UUCGCG) has a 5’-UU-overhang exhibiting Hoogsteen-like trans U.U base pairs. Nat Struct Biol, 3, 24–31.

33. Bartoszewski, R. and Sikorski, A.F. (2019) Editorial focus: understanding off-target effects as the key to successful RNAi therapy. Cell Mol Biol Lett, 24, 69.

34. Elbashir, S.M., Martinez, J., Patkaniowska, A., Lendeckel, W. and Tuschl, T. (2001) Functional anatomy of siRNAs for mediating efficient RNAi in Drosophila melanogaster embryo lysate. EMBO J, 20, 6877–6888.

35. Hadjadj, J., Yatim, N., Barnabei, L., Corneau, A., Boussier, J., Smith, N., Pere, H., Charbit, B., Bondet, V., Chenevier-Gobeaux, C. et al. (2020) Impaired type I interferon activity and inflammatory responses in severe COVID-19 patients. Science, 369, 718–724.

36. Broggi, A., Ghosh, S., Sposito, B., Spreafico, R., Balzarini, F., Lo Cascio, A., Clementi, N., De Santis, M., Mancini, N., Granucci, F. et al. (2020) Type III interferons disrupt the lung epithelial barrier upon viral recognition. Science.

37. Park, A. and Iwasaki, A. (2020) Type I and Type III Interferons - Induction, Signaling, Evasion, and Application to Combat COVID-19. Cell Host Microbe, 27, 870–878.

38. Wadman, M. (2020) Can interferons stop COVID-19 before it takes hold? Science, 369, 125–126.

39. Wadman, M. (2020) Can boosting interferons, the body’s frontline virus fighters, beat COVID-19? Science.

40. Lokugamage, K.G., Hage, A., Schindewolf, C., Rajsbaum, R., and Menachery, V.D. (2020) SARS-CoV-2 is sensitive to type I interferon pretreatment. bioRxiv.

41. Mantlo, E., Bukreyeva, N., Maruyama, J., Paessler, S. and Huang, C. (2020) Antiviral activities of type I interferons to SARS-CoV-2 infection. Antiviral Res, 179, 104811.

42. Hung, I.F., Lung, K.C., Tso, E.Y., Liu, R., Chung, T.W., Chu, M.Y., Ng, Y.Y., Lo, J., Chan, J., Tam, A.R. et al. (2020) Triple combination of interferon beta-1b, lopinavir-ritonavir, and ribavirin in the treatment of patients admitted to hospital with COVID-19: an open-label, randomised, phase 2 trial. Lancet, 395, 1695–1704.

43. Thoms, M., Buschauer, R., Ameismeier, M., Koepke, L., Denk, T., Hirschenberger, M., Kratzat, H., Hayn, M., Mackens-Kiani, T., Cheng, J. et al. (2020) Structural basis for translational shutdown and immune evasion by the Nsp1 protein of SARS-CoV-2. Science, 369, 1249–1255.

44. Jain, A., Barrile, R., van der Meer, A.D., Mammoto, A., Mammoto, T., De Ceunynck, K., Aisiku, O., Otieno, M.A., Louden, C.S., Hamilton, G.A. et al. (2018) Primary Human Lung Alveolus-on-a-chip Model of Intravascular Thrombosis for Assessment of Therapeutics. Clin Pharmacol Ther, 103, 332–340.

45. Huh, D., Leslie, D.C., Matthews, B.D., Fraser, J.P., Jurek, S., Hamilton, G.A., Thorneloe, K.S., McAlexander, M.A. and Ingber, D.E. (2012) A human disease model of drug toxicity-induced pulmonary edema in a lung-on-a-chip microdevice. Sci Transl Med, 4, 159ra147.

46. Huang da, W., Sherman, B.T. and Lempicki, R.A. (2009) Systematic and integrative analysis of large gene lists using DAVID bioinformatics resources. Nat Protoc, 4, 44–57.

47. Lewis, K.B.a.S.R.a.M. (2020) EnhancedVolcano: Publication-ready volcano plots with enhanced colouring and labeling.

48. Navarrete-Perea, J., Yu, Q., Gygi, S.P. and Paulo, J.A. (2018) Streamlined Tandem Mass Tag (SL-TMT) Protocol: An Efficient Strategy for Quantitative (Phospho)proteome Profiling Using Tandem Mass Tag-Synchronous Precursor Selection-MS3. J Proteome Res, 17, 2226–2236.

49. Gygi, J.P., Yu, Q., Navarrete-Perea, J., Rad, R., Gygi, S.P. and Paulo, J.A. (2019) Web-Based Search Tool for Visualizing Instrument Performance Using the Triple Knockout (TKO) Proteome Standard. J Proteome Res, 18, 687–693.

50. Paulo, J.A., O’Connell, J.D. and Gygi, S.P. (2016) A Triple Knockout (TKO) Proteomics Standard for Diagnosing Ion Interference in Isobaric Labeling Experiments. J Am Soc Mass Spectrom, 27, 1620–1625.

51. Huttlin, E.L., Jedrychowski, M.P., Elias, J.E., Goswami, T., Rad, R., Beausoleil, S.A., Villen, J., Haas, W., Sowa, M.E. and Gygi, S.P. (2010) A tissue-specific atlas of mouse protein phosphorylation and expression. Cell, 143, 1174–1189.

52. Elias, J.E. and Gygi, S.P. (2010) Target-decoy search strategy for mass spectrometry-based proteomics. Methods Mol Biol, 604, 55–71.

53. Elias, J.E. and Gygi, S.P. (2007) Target-decoy search strategy for increased confidence in large-scale protein identifications by mass spectrometry. Nat Methods, 4, 207–214.

54. McAlister, G.C., Huttlin, E.L., Haas, W., Ting, L., Jedrychowski, M.P., Rogers, J.C., Kuhn, K., Pike, I., Grothe, R.A., Blethrow, J.D. et al. (2012) Increasing the multiplexing capacity of TMTs using reporter ion isotopologues with isobaric masses. Anal Chem, 84, 7469–7478.

